# A comprehensive atlas of immunological differences between humans, mice and non-human primates

**DOI:** 10.1101/574160

**Authors:** Zachary B Bjornson-Hooper, Gabriela K Fragiadakis, Matthew H Spitzer, Deepthi Madhireddy, Dave McIlwain, Garry P Nolan

## Abstract

Animal models are an integral part of the drug development and evaluation process. However, they are unsurprisingly imperfect reflections of humans, and the extent and nature of many immunological differences are unknown. With the rise of targeted and biological therapeutics, it is increasingly important that we understand the molecular differences in immunological behavior of humans and model organisms. Thus, we profiled a large number of healthy humans, along with three of the model organisms most similar to humans: rhesus and cynomolgus macaques and African green monkeys; and the most widely used mammalian model: mice. Using cross-species, universal phenotyping and signaling panels, we measured immune cell signaling responses to an array of 15 stimuli using CyTOF mass cytometry. We found numerous instances of different cellular phenotypes and immune signaling events occurring within and between species with likely effects on evaluation of therapeutics, and detail three examples (double-positive T cell frequency and signaling; granulocyte response to *Bacillus anthracis* antigen; and B cell subsets). We also explore the correlation of herpes simian B virus serostatus on the immune profile. The full dataset is available online at https://flowrepository.org (accession FR-FCM-Z2ZY) and https://immuneatlas.org.

## Introduction

Animal models are core to drug development and evaluation: all candidate therapeutics are evaluated for toxicity, pharmacokinetics/pharmacodynamics (PK/PD) and efficacy using animals. Furthermore, in circumstances when human trials of efficacy cannot be conducted for ethical reasons or when an insufficient number of natural cases exists, such as for therapeutics for biothreat agents, the U.S. Food and Drug Administration may allow licensure after efficacy is evaluated only in animal models under 21 CFR 314 (“The Animal Rule”). This rule has been applied in a limited number of cases since its issuance in 2002, including Raxibacumab for inhalational anthrax, B12 for cyanide poisoning, pyridostigmine for nerve gas and levofloxacin and moxifloxacin for bubonic plague^1^, but therapeutics for an increasing number of conditions are being evaluated via this mode.

Unsurprisingly, animal models are not perfect surrogates for humans (see Davis *et al*.^2^, among other reviews). At a wide view, most diseases infect only a limited number of species. Modeling these diseases is inherently challenging, and researchers use a variety of imperfect, compensatory techniques. For example, naturally occurring ebolaviruses do not infect mice, so researchers use mouse-adapted viruses, which still do not produce disease very similar to that seen in humans^3^. Cancers are slow to induce and rarely develop naturally over the short lifetime of a mouse, so tumors are xenografted into immunocompromised mice or an artificially small number of genes are mutated to induce tumor formation^4^.

Furthermore, even when a model appears reflective of a disease at a wide view, *e.g*. by physical examination, differences in finer details such as cellular receptors or kinases may adversely affect evaluation of therapeutics. Broadly characterized animal models may have sufficed for evaluation of the classical vaccines, antibiotics and other broadly active and agent-targeting therapeutics that were the focus of medicine for the century since Pasteur used sheep to model anthrax in the 1880s. However, understanding the molecular and cellular differences in immune signaling between animal models and humans is of increasing importance as more rationally designed biotherapeutics that target the host, and especially specific receptors and pathways, are brought to trial. For example, on a molecular level, while human CD16 (the type III Fcγ receptor) interacts with IgG1 and IgG3, macaque CD16 instead interacts with IgG1 and IgG2^5^; additionally, CD16 is absent from macaque granulocytes^5,6^. These differences will likely confound evaluation of therapeutic antibodies that act through this Fcγ receptor^7^. On a systems level, there is almost no correlation of transcriptomic responses to burn, trauma and endotoxemia between humans and mice^8^, underscoring that these species have evolved unique mechanisms to heal and combat disease. While expression patterns of some orthologous genes may be generally similar between species, a large number of genes have divergent expression patterns; regulatory elements especially have a lower level of conservation across species, as do lineage-specific responses^9,10^.

In addition to being imperfect efficacy models, animals are also imperfect safety models. For example, five people died in Fialuridine clinical trials when the drug was administered at doses that were nontoxic in mice, rats, dogs and monkeys^11^—four very different species. TGN1412 (anti-CD28) caused catastrophic organ failure in humans when administered at 1/500^th^ of the safe animal dose due to differences in CD28 expression in the model organisms used^12^. The overall low accuracy of animal models also begs the question *how many drugs would work in humans but fail in our animal models?*

Finally, it is also important to consider differences between human demographic groups used in clinical trials in terms of ages, genders, ethnicities, immunological histories, genetics, etc. These differences are perhaps better recognized than those between animal models and humans: the US Food and Drug Administration since 1998 has required that new drug applications describe safety and effectiveness by gender, age and race, and the labels for 38% of drugs approved from 2004 to 2007 include PK/PD data by ethnicity^13^. Warfarin, rosuvastatin, tacrolimus, carbamazepine and others are known to have ethnicity-dependent PK/PD^13^. A variety of factors, including hormones, body fat and blood flow, contribute to gender differences in bioavailability, distribution, metabolism and excretion of drugs^14^.

To address these issues and systematically characterize the immunological variability both within and between species, we performed phospho-flow immune signaling profiling of whole blood from 86 healthy humans, 32 rhesus macaques (*Macaca mulatta*), 32 cynomolgus macaques (*Macaca fascicularis*), 24 African green monkeys (*Chlorocebus aethiops*) and 50 C57BL/6 mice (*Mus musculus*), measuring 16 signaling proteins and 24 surface markers per cell in approximately 20 immune cell populations after treatment with a panel of 15 stimuli, using CyTOF mass cytometry. The set of phenotyping panels was designed to demarcate orthologous populations in all species, while the signaling panel and complementary stimuli were targeted at innate immunity and cytokine responses. We present an overview of this cross-species analysis here, including several examples of specific differences between species. The complementary manuscript by Fragiadakis *et al*.† analyzes the human dataset in detail, including an examination of demographic differences, presenting a reference for human immune profiling studies. The entirety of the dataset is available in an online analysis portal at https://immuneatlas.org.

## Results

### Standardization and Validation of Phenotyping, Signaling and Stimuli Assays

Our goal was to create a high-quality dataset with minimal technical variability. The first step toward this was to create universal antibody panels that allowed parallel investigation of cell phenotypes and behaviors across species. Previously we reported the design and validation of a universal CyTOF phenotyping panel for humans and three species of non-human primates that delineates the same populations in all species while using the same antibody epitopes and clones whenever possible to minimize technical bias (Bjornson-Hooper *et al.*, in preparation). This panel was based on a screen of more than 300 commercially available antibodies as well as thorough literature review and targeted experiments. Here we created a similar panel for mice that delineates the same populations as those targeted by the primate panel (Table S1) and enables parallel gating of all five species (Figure S1). Note that these hierarchies only show the major populations and make use of a subset of our phenotyping panel. It is possible to further subset most of these populations—for example, NK cells can be further subset based on CD20, CCR7 and CD56—and it is our hope that interested researchers will explore the raw data on https://flowrepository.org (accession FR-FCM-Z2ZY) or https://immuneatlas.org.

We designed the stimuli panel and complementary signaling antibody panels in tandem. Signaling epitopes are highly conserved and, with only one exception, the same antibody clones were usable in all five species (see Methods). The stimuli and readouts were selected for their clinical relevance and their roles in disease and innate immunity (see Methods). Many of the stimuli are recombinant cytokines; these recombinant proteins were preferably species-matched; *i.e*. recombinant rhesus cytokines were used in rhesus macaque blood. Where no suitable proteins were found, we used the closest species available (*e.g*. macaque cytokines were frequently used in African green monkey blood).

Many steps were taken to reduce error and technical variability (see Methods). Antibodies were conjugated in bulk and then cryo-lyophilized into single-use cocktail pellets for stability and to eliminate pipetting errors from repeatedly assembling cocktails. Stimuli were pre-dispensed into single-use plates. Fresh, whole blood was used instead of PBMCs to avoid perturbations and inconsistency from density gradient isolations and storage. We used mass-tag barcoding to combine staining of all stimulation conditions into one tube per donor, eliminating differences in staining volume and processing. Complete automation was used for stimulation, fixation, lysis, barcoding and staining. CyTOF quality control tests were run before every sample, and internal normalization standards were included to control for variability during runs. Finally, all analysis code has been annotated and made available, along with the detailed records from antibody conjugations and sample processing.

### Cell type frequencies vary between species and recapitulate the evolutionary tree

Using the hierarchy shown in Figure S1, we gated blood cells from all species in parallel to determine the distributions of frequencies of these cell types (Figure 1). The frequencies of many populations were significantly different between species; notable differences include (a) macaques have more CD4+/CD8+ double-positive T cells than humans, mice or AGMs, (b) mice have approximately 10 times fewer neutrophils than all primates, (c) all non-human primates have approximately three times more B cells than humans, and mice have approximately 10 times more than humans, and (d) humans have a higher ratio of classical to non-classical monocytes than any other species. Due to space constraints, only the first finding is discussed in detail here; the rest can be viewed at https://immuneatlas.org.

**Figure 1.**
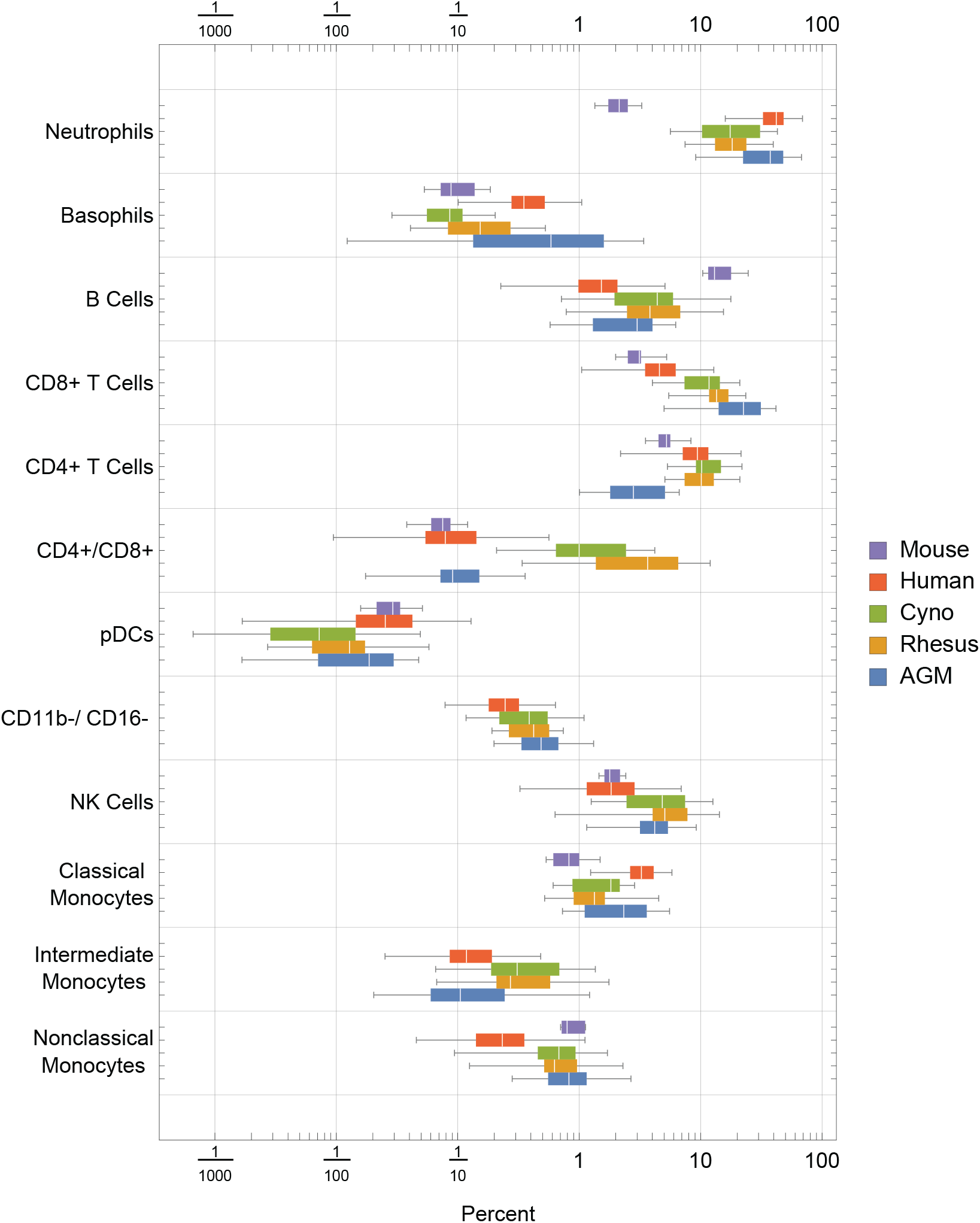
Frequencies (percent of total) of gated cell types by species. Center line: median; Box: 25th to 75th quantile; Whiskers: 1.5x interquartile range.

When we clustered the species by their average cell type frequencies, we recapitulated the evolutionary tree (Figure 2, right panel): rhesus and cynomolgus macaques diverged most recently, 1.5 to 3.5 million years ago (Ma); macaques and African green monkeys diverged 11.5 to 14 Ma; Old World monkeys and humans diverged 20-38 Ma and mice diverged more than 90 Ma^15–17^. This result raises an intriguing possibility for future analysis of a large number of species to look for critical junctures in the evolution of the immune system.

**Figure 2.**
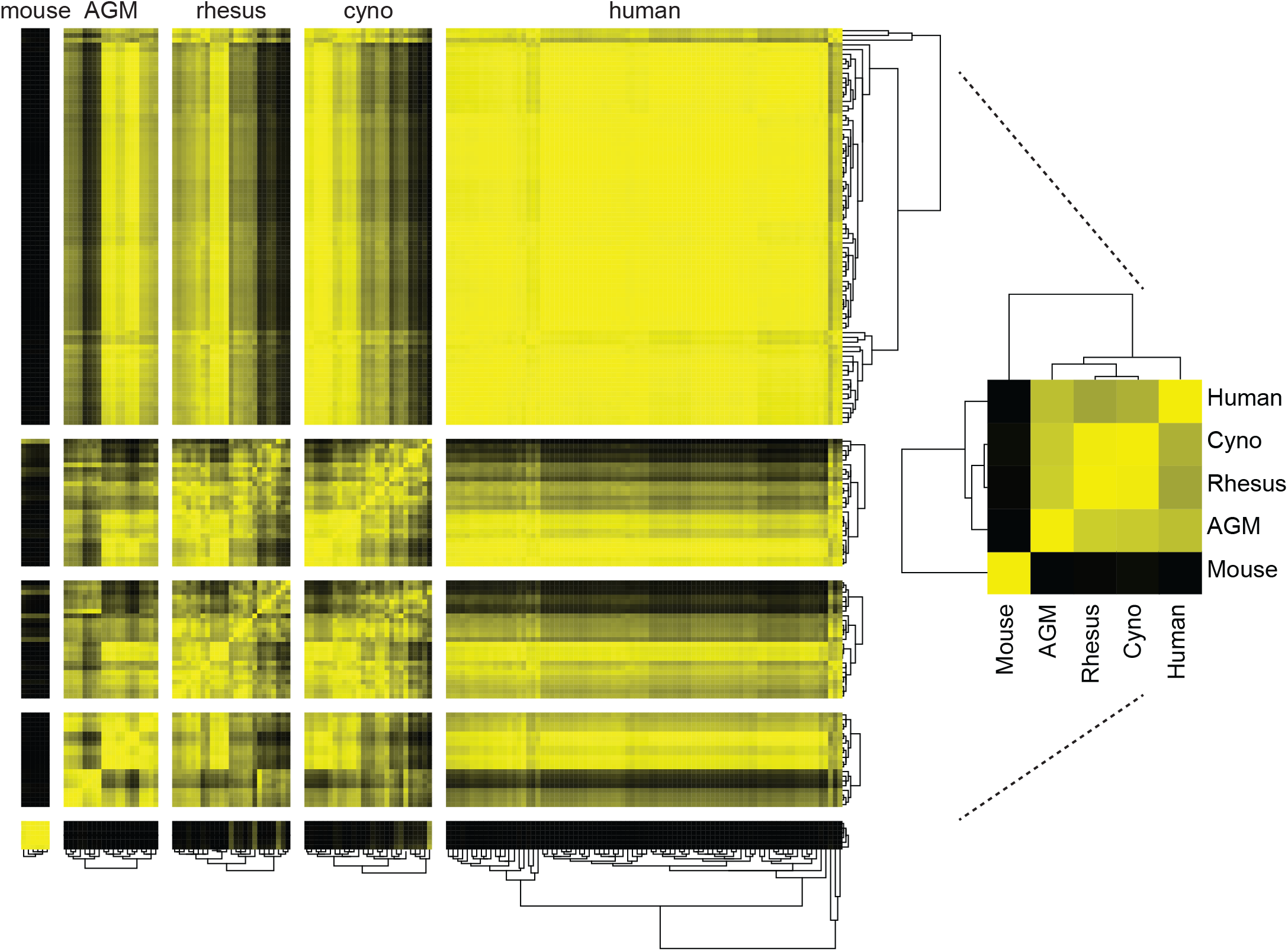
Clustering of species by their cell type frequencies recapitulates the evolutionary tree. The average frequencies of 10 cell types in each species was calculated, and then clustered according to their pairwise correlations between species (right). This major clustering order was preserved in the larger heatmap (left), in which each individual donor is displayed and clustered by their individual cell type frequencies. Metric: PearsonCorrelation[freqs_species_1, freqs_species_2]^2^; distance function: Euclidean; linkage: average.

While maintaining this major clustering order, we then sub-clustered donors within their species, again by their cell type frequencies (Figure 2, left panel). The intra-species distances between clusters were the smallest for mice, followed by the non-human primates, followed by humans. This is unsurprising considering that the mice were a single inbred strain, the non-human primates were either wild-caught from isolated colonies or captive-bred, and the humans were specifically recruited to include a variety of ethnicities. For the most part, all individuals within a species had similar correlations to other species; nonetheless, some non-human primates were more similar to humans than others. Within the humans, clustering did not strongly correlate with age, gender or ethnicity. Importantly, clustering did not correlate with the batches in which individuals were processed; that is, batch effects are not a driver of this clustering.

### Orthologous cell types in different species have unique phenotypes

To assess differences in phenotypes of the orthologous cell populations between species, we plotted the distributions of staining intensities of every surface marker in every species by population (example in Figure 3, full set in Figure S2). While these populations express many of the same markers across species, we found many significant differences across a wide range of cell types. For example, neutrophils in humans express high levels of CD16 and moderate levels of CD11c, in contrast to all three NHP species; meanwhile, neutrophils in the three NHP species express higher levels of CCR7 and neutrophils in mice express moderate levels of CD16/32. B cells in macaques express higher levels of CD1c than in humans or African green monkeys, as discussed later. CD4+ and CD8+ T cells in humans express higher levels of CD161 and CD7 than NHPs, while mouse T cells do not express NK1.1 (CD161). NK cells in NHPs express higher levels of CD8 than humans; this is consistent with Autissier *et al*.,^18^ although one must not ignore the fact that some human NK cells also express CD8^19^. Classical monocytes in African green monkeys stain brightly for BDCA3—a canonical dendritic cell subset marker in humans.

**Figure 3.**
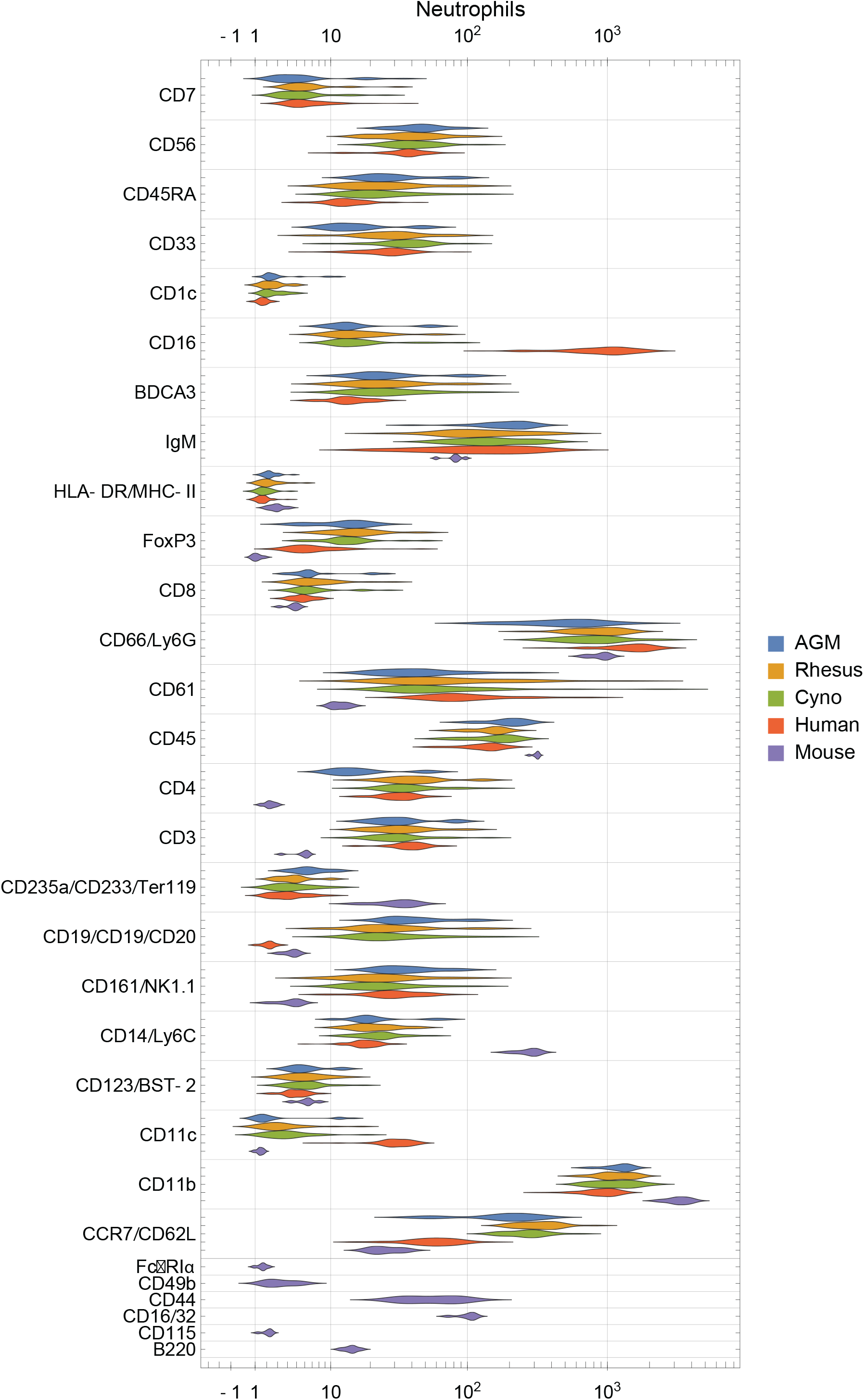
Distribution of surface marker expression for each species in neutrophils. (See Figure S2 for other populations.) Different markers are grouped together if they were on the same channel and stain similar cell types between species (e.g. CD235a/CD233/Ter119 are all on In113 and stain erythrocytes), and are labeled as “[all species]”, “[primates]/[mice]” or “[humans]/[non-human primates]/[mice]”.

Although these particular differences are almost certainly biological, differences in staining intensities should be interpreted with caution because genetic variation of antigens may affect binding affinity between species. One must also consider the possibility for cross-reactivity with different epitopes and populations in NHPs than in humans, against which most of the primate panel antibodies were raised. For this reason, we primarily focus on cases where a marker is present in one or more species and essentially absent in another, or cases that we can corroborate through further data mining and literature review.

### Species-specific signaling profiles have profound implications for drug development

To study functional differences of orthologous cell types, we visualized signaling behaviors by species (Figure 4 and S3). These charts are meant to provide a compact, semi-quantitative overview of this large dataset. While we are unable to explore and validate every one of the many differences in detail, we explore several briefly here, and several in depth in the following sections.

**Figure 4.**
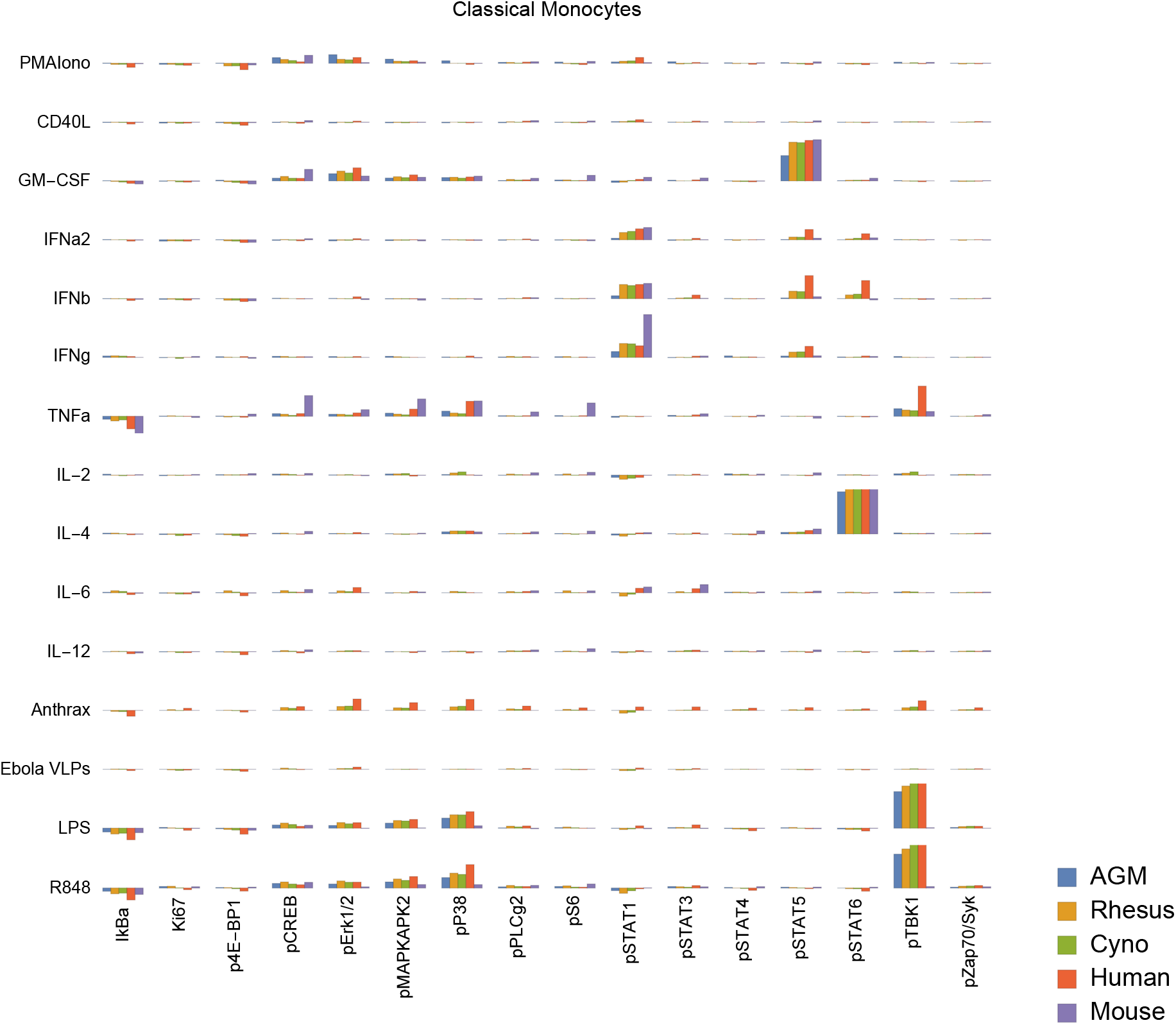
Signaling responses (difference of ArcSinh-transformed values, or approximately fold-change) in classical monocytes by stimulus, activation marker and species (other cell types in Figure S3). Note that Bacillus anthracis (“anthrax”) and Ebola VLPs were not available for use as stimuli in mice or AGMs; thus, values for these species are always displayed as zero. We could not gate intermediate monocytes or a “CD11b-/CD16-”-equivalent population in mice; these values are also zero. The Y axis range of all charts is −0.5 to +1.5.

In terms of innate immune responses, we observed an absence of TLR7/8 (R848) and TLR4 (LPS) responses in mouse classical monocytes, as evidenced by a lack of pTBK1, pP38 and pMAPKAPK2 signaling. The TLR7/8 (R848) results are not unexpected since prior literature indicates that mouse TLR8 is defective^20^, and TLR7 is primarily expressed in DCs^21^ (and, consistently, we saw a response to R848 in mouse pDCs). The lack of TLR4 signaling in mouse monocytes is potentially novel; expression and functionality of TLR4 in mouse blood monocytes under basal conditions is unclear from prior literature, but are known to have significant differences from humans^22,23^. Considering these findings in addition to the fact that mouse monocytes essentially lack MHC-II and are thus not major antigen-presenting cells^24,25^, it is clear that classical monocytes, which are crucial to innate immune responses in humans, serve a very different role in mice.

We also observed that all three NHP species have an almost-absent pSTAT6 response to IL-4 in non-classical monocytes. This is in sharp contrast to humans and mice, where pSTAT6 responses to IL-4 are a major, canonical signaling response. Mice deficient in STAT6 exhibit defective immune behaviors in response to IL-4 across the spectrum^26,27^: B cells do not upregulate MHC-II or CD23 and do not proliferate when co-stimulated with anti-IgM; T cells show reduced proliferation in response to PMA; T cell cytokine production and allergic antibody responses are reduced; T cells also fail to differentiate into Th2 cells. Despite the almost absent response in non-classical monocytes, all three NHP species have varying degrees of pSTAT6 responses to IL-4 in other cell types, including T cells, classical monocytes, intermediate monocytes, neutrophils, B cells, DCs and NK cells. These responses serve as positive technical controls for STAT6 and IL-4 in NHPs in our assays, but may also point towards biological compensation for the lack of response in non-classical monocytes in NHPs.

Finally, we observed that African green monkey pDCs have essentially no IkBα response to R848, although they have a stronger pTBK1 response than any of the other species. Meanwhile, cynomolgus pDCs have a unique pSTAT5 response to IL-6. Others have reported that IL-6 causes low-levels of phosphorylation of STAT5 in T cells and NK cells in mice to detectable levels by 15 minutes^28^ (the time point we used); we too see a small response in mice and humans in those cell types, but the magnitude of these responses are dwarfed by that of cynomolgus pDCs.

These results highlight the importance of careful selection and interpretation of animal models for drug development and evaluation, as the substantial signaling differences between species could provide misleading results in terms of drug responsiveness at the cellular level. The remainder of this report details several specific examples of differences between species in B cells, T cells, granulocytes and monocytes, particularly focusing on several cell types and behaviors unique to macaques.

### Macaque monocyte abundance correlates with herpes simian B virus status

Many macaques are infected with herpes simian B virus (formerly *Cercopithecine herpes virus 1*, or B virus) by natural exposure. Like herpes simplex viruses in humans, the virus is community- or sexually acquired, persists for life and is essentially harmless to macaques (When transmitted to humans, the disease is usually lethal.) African green monkeys are not natural carriers of herpes simian B virus, although they are carriers of another herpes virus, simian agent 8 (SA8, *Cercopithecine herpes virus 2*)^29^. Whether or not herpes serostatus has an effect on general immunological health is unknown, and as far as we are aware, no published study in macaques has ever taken into consideration the serostatus of the animals.

We detected a significant association between B virus status and non-classical monocyte abundance in all macaques (Figure 5a), and a stronger association with both intermediate and non-classical monocyte frequency in rhesus macaques (Figure 5b). This finding suggests that herpes simian B virus serostatus can indeed affect immune function and may thus factor into innate immune responses observed during therapeutic evaluations performed in macaques. Future work should segregate animals with latent and active infection, which we hypothesize will yield an even stronger correlation with immunological factors.

**Figure 5.**
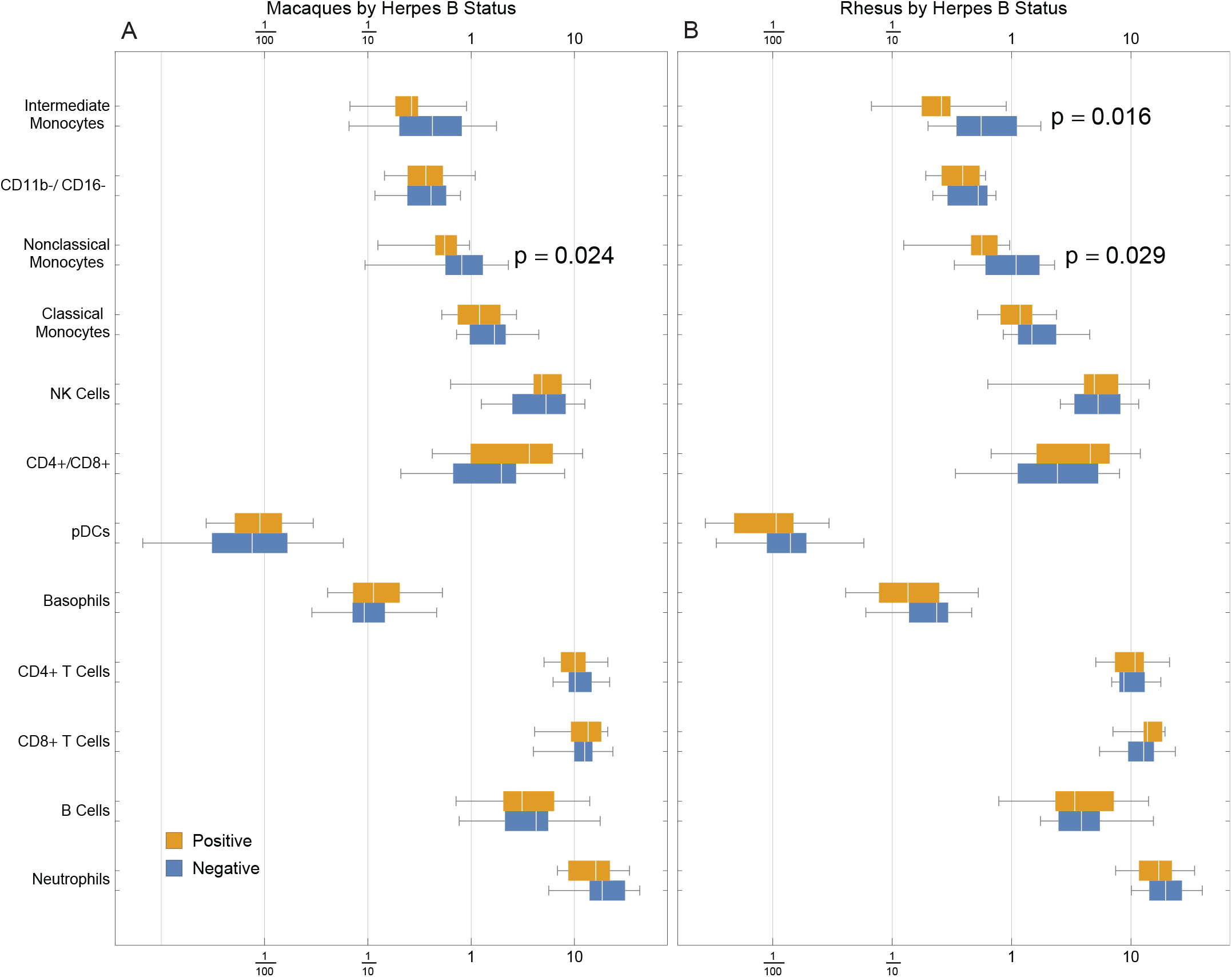
Frequencies of cell types in rhesus and cynomolgus macaques (a) or rhesus macaques only (b) by herpes simian B virus status.

### CD4+CD8+ double-positive T cells are more abundant in macaques and display unique responses to different type I interferons

Macaques are known to have higher frequencies of peripheral CD4+CD8+ double-positive (DP) T cells than humans, the frequency of which furthermore increases with age (Table S2). We found a median frequency of 5.3% in rhesus, 1.4% in cynomolgus and less than 0.2% in AGM, mouse and human (Figure S4). While in humans DP T cells were initially considered immature cells that were prematurely released from the thymus, more recent work has found the population to contain differentiated, effector-memory lymphocytes^30^. In macaques, DP T cells appear to be a mature, normal population^31,32^ (reviewed in^33^). These cells are also associated with viral infections in humans^34^, mice^35^ and chimpanzees^30^, and are found in the intestine in mice^36^.

Excitingly, we observed that macaque DP T cells respond differently to IFNα and IFNβ (Figure S5). While all type 1 interferons signal through the same receptor complex^37^ and are thus expected to have largely redundant downstream signaling effects, there are a number of known differences: for example, IFNβ is more potent than IFNα in inducing apoptotic and anti-proliferative pathways, and IFNβ alleviates some effects of multiple sclerosis while IFNα does not (see^38^ for a review of differences). The mechanisms of many of these differences are unknown. We found that macaque double-positive T cells have a strong pSTAT5 response to IFNβ but not to IFNα (Figure S5); meanwhile, the pSTAT1 and pSTAT4 responses to both IFNs followed the same trends across T cell subsets.

### Macaque granulocytes are rapidly responsive to *Bacillus anthracis*

We evaluated the signaling response to a 15-minute incubation with 22 × 10^6^ CFU of gamma-irradiated (inactivated), vegetative *Bacillus anthracis* Ames. Despite the short incubation period, we found a subtle, but significantly greater, level of neutrophil activation (Ki67) in macaques than in humans (Figure 6). This indicates a very rapid response to infection in these animals; humans could lack this response outright or have delayed kinetics.

**Figure 6.**
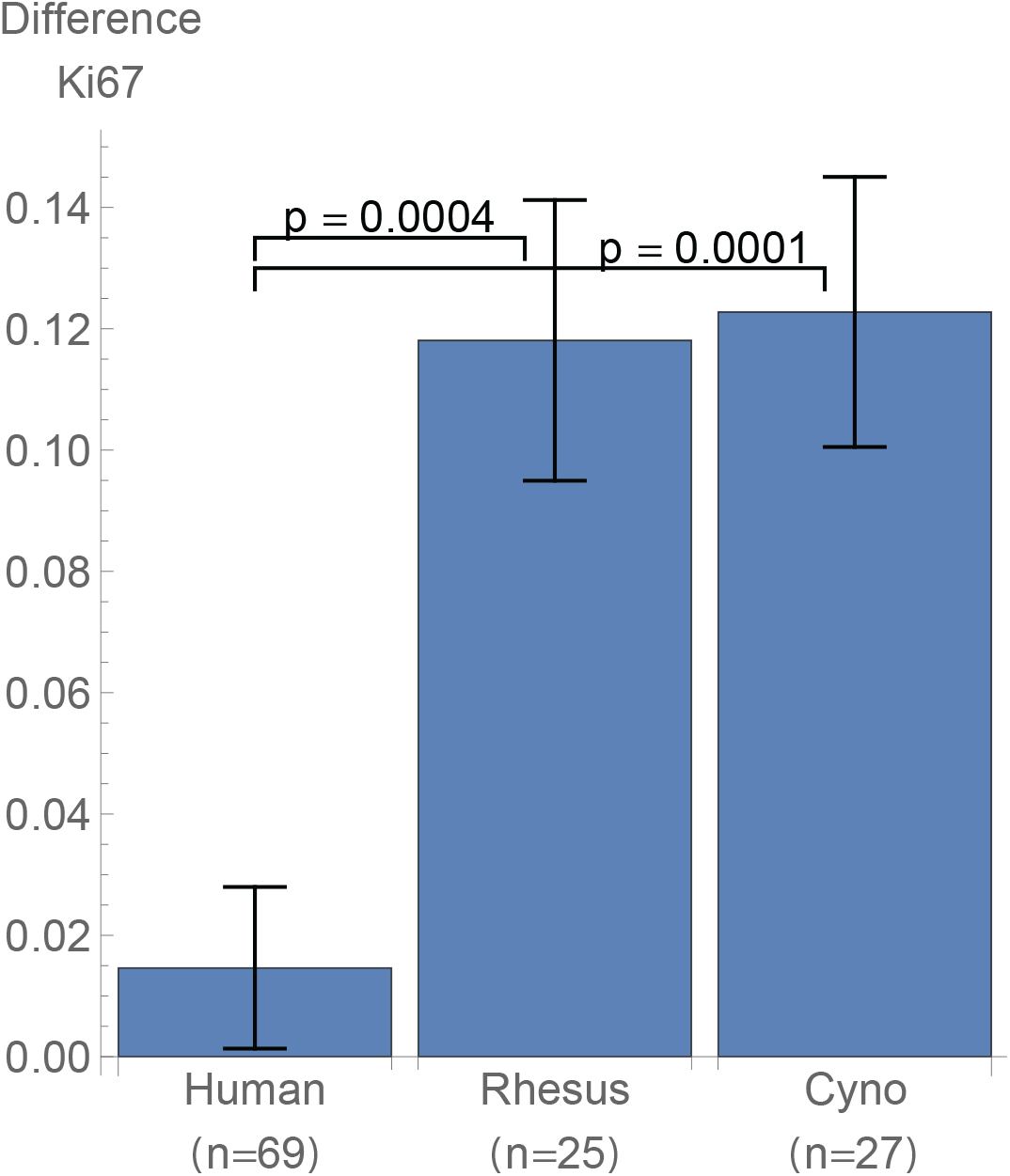
Neutrophil Ki67 induction (mean and standard error of differences of ArcSinh-transformed values) by species after exposure to 22M CFU of gamma-irradiated Bacillus anthracis for 15 minutes.

That this activation occurs in neutrophils is especially interesting not only because neutrophils kill *B. anthracis*^39^, but also because macaque and several other NHP species’ neutrophils uniquely contain theta defensins^40^, which are highly potent antibiotics against *B. anthracis* and its lethal factor (LF)^41,42^. This finding is particularly important considering that anthrax therapeutics are among those that are candidates for evaluation under The Animal Rule, with efficacy studies conducted in animal models.

For reasons beyond our control^43^, we discontinued use of *Bacillus* antigen midway through the project; thus, none of the mouse or AGM samples were treated with *Bacillus* antigen, nor were 18 of the human samples. Accordingly, our analyses considered only the samples that were treated.

### Macaques have unique CD1c+ and CD8+ B cell subsets

We observed two unique subsets of B cells in macaques defined by either CD1c or CD8α expression. These populations were either rare or absent in humans, mice and African green monkeys, and their presence and donor-to-donor variability likely has implications for disease modeling.

CD1 is a family of lipid and glycolipid-presenting molecules—a counterpart to MHC class I and II found on a subset of B cells and dendritic cells that plays an important role in humans in defense against diseases such as tuberculosis. CD1c (BDCA-1) specifically presents mannosyl mycoketide and phosphomycoketide^44^. A previous study reported that 21.4% of B cells in rhesus macaques were CD1c+ B cells, in contrast to humans with only 3.3%^18^. We similarly found significantly more CD1c+ B cells in all three NHP species that we examined when compared to humans, and significantly higher amounts of CD1c therein (Figure 7a and b). Interestingly, mice and rats lack group 1 CD1 altogether^44^. Mouse susceptibility to tuberculosis varies by strain, but at least several, including the common C57BL/6 and BALB/c strains, are resistant^45^, and thus must have group 1 CD1-independent mechanisms for controlling infection. With regard to the higher intensity of CD1c staining, we initially considered that this could be due to a difference in antibody affinity between species; however, the levels of CD1c on DCs were similar between macaques and humans (Figure 7c). Considering that the higher abundance of CD1c would likely correspond to a larger display of mycobacterial antigens, this difference could indicate that macaques have an increased ability to invoke an adaptive response to mycobacterial infections. Alternatively, higher CD1c abundance could be compensating for a weaker related part of the immune system; for example, they could have fewer or less-responsive CD1-restricted T cells and/or less effective upstream lipid processing, such as the deglycosylation of mycobacterial mannosyl phosphomycoketide.

**Figure 7.**
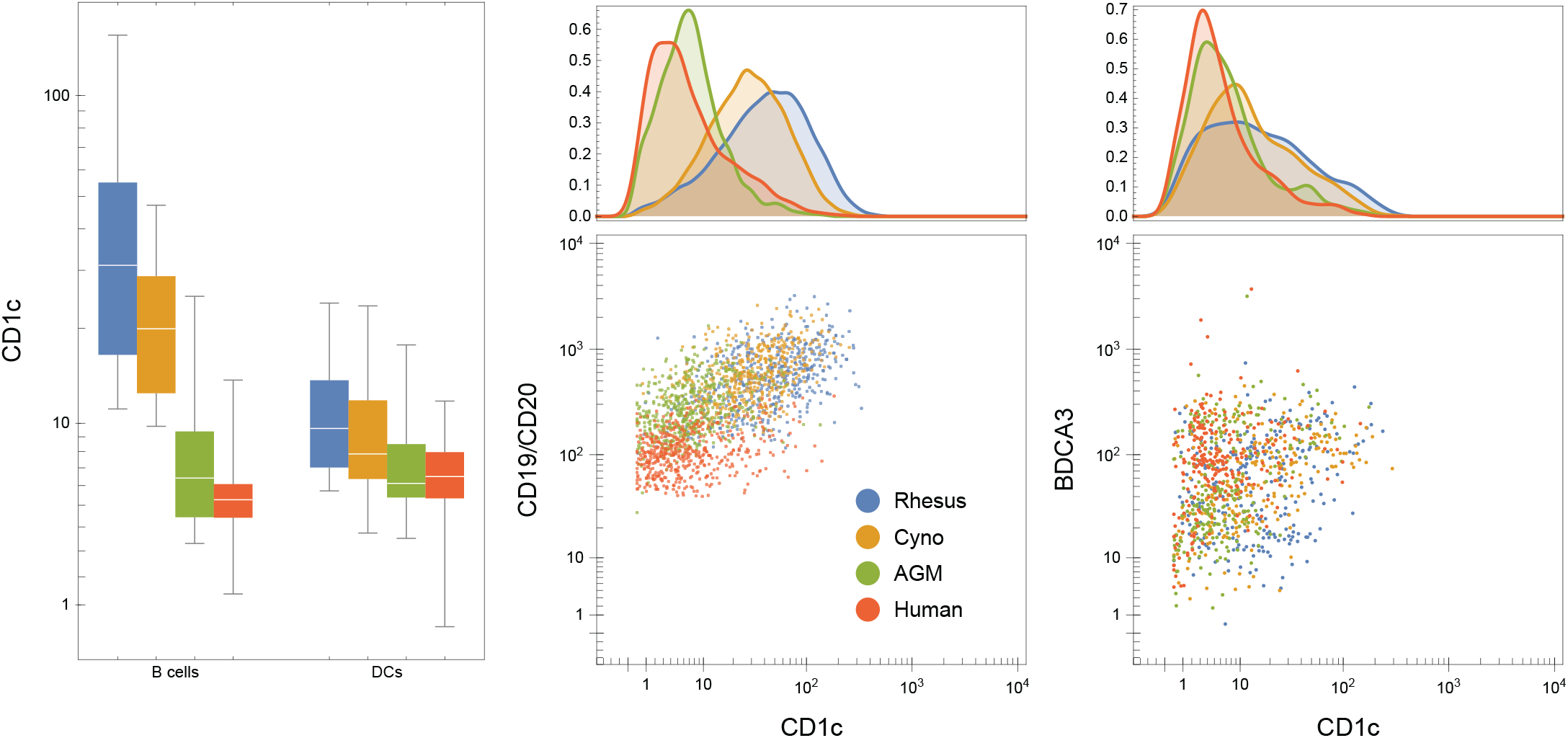
CD1c+ B cells are more abundant in non-human primates than in humans, and CD1c is furthermore expressed at higher levels in NHP B cells, especially in macaques. Left: medians CD1c staining of all individual donors by species. Middle and right: One representative individual from each species. Dot plots show 500 randomly selected B cells (middle) or 250 randomly selected CD11b-/CD16-DCs (right).

In humans, CD8α is found almost exclusively on T cell and NK cell subsets; exceptions are limited to conditions such as HIV-1^46^, B-cell leukemia^47–49^ and lymphomas^50^, and potentially a very small subset in healthy individuals^46^. In many rhesus macaques, however, CD8α is also found on a subset of B cells^51^. We found 19 out of 25 animals had at least 0.5% of their B cells stain for CD8 and some animals had as much as 33% of B cells stain (mean: 6.5%, median: 3.3%) (Figure S6). We gated these cells as CD45+ CD66-CD3-CD20+ CD7-CD8+, thereby excluding T cells and NK cells, which also stain for CD8 in macaques. We continued to determine that cynomolgus macaques, African green monkeys and mice do not appear to have CD8+ B cells (Figure S7). Because we simultaneously measured phenotyping and signaling molecules, we were also able to evaluate the functional behavior of CD8+ B cells (Figure S8). Considering several higher-magnitude responses, we found that these cells still display the hallmark signaling behaviors of CD8-B cells, albeit perhaps with stronger pSTAT1 signaling.

These findings are thus potentially important criteria for selecting not only which *species* to use for model development and therapeutic evaluation, but given the high variance of CD8+ B cell frequency within rhesus macaques, which *individual donors*.

### Public Web portal for further exploration

Ultimately, we generated nearly 3,000 flow cytometry standard (FCS) files containing more than one billion cellular events. To allow easy access to the data by other researchers, we posted the curated dataset along with animal health reports, human donor questionnaires and *Mathematica* notebooks used to create the figures in this manuscript to https://flowrepository.org (accession FR-FCM-Z2ZY) and https://immuneatlas.org. All protocol records and robotics logs are available upon request.

## Discussion

Animal models are commonplace in drug development but are a challenging part of the process. Research based on imperfect animal models is wasteful through both false positives, which work in models but fail in humans, and false negatives, which fail to work in models but would work in humans. Aside from the economic and ethical burden, this creates a public health problem by diminishing researchers’ abilities to develop therapeutics and countermeasures for emerging diseases.

Here we show that model species must be carefully selected and evaluated based on their relevance to the particular experiment at hand. For example, a therapeutic targeting one pathway in one cell type should be confirmed to trigger relevant, similar signaling in humans as in the model species. Nonetheless, even with careful selection based on evaluation of one or several particular pathways, one cannot discount the complex interplay between multiple pathways and multiple cell types. In some cases, highly targeted immunotherapeutics seem nearly impossible to validate in animal models because of this.

One might counter that for over a century we have used animal models for development of many successful vaccines and drugs. However, some of the most successful therapeutics, such as the smallpox vaccine, are broadly immunogenic and activate large swaths of the immune system. Broad activation is likely to be inherently less susceptible to minute differences in cellular immunology between species. Similarly, antibiotics—perhaps the most important class of therapeutics in history—target the infectious agent instead of host processes; thus, efficacy is far less dependent on the host species. Successfully developing the next generation of precision targeted therapeutics acting on the host will require careful selection of relevant models.

For these reasons, researchers should continue to consider humans as early as possible in the drug development process, including in initial planning, screening and evaluation stages. For example, using primary human cells (*e.g*. blood and tumor samples) in place of cell lines and cells from other species, as well as evaluating differences in mechanisms of disease and therapeutics between humans and model organisms will likely yield dozens of therapeutic candidates that have been missed because they have been evaluated in models that do not accurately reflect human immunology for a sufficient evaluation of that candidate.

In the analysis of cell phenotypes presented here, we focused on high-confidence differences between species. There are several observed differences that we did not discuss for specific reasons: (a) While human NK cells, B cells and non-classical monocytes appear to express higher amounts of CD45RA, we previously observed what could be a difference in affinity for this clone between NHPs and humans. (b) Because humans were stained with anti-CD19 and NHPs with anti-CD20, we cannot necessarily conclude that the difference in staining of these B cell markers is significantly different. (c) As discussed earlier, although CD11c is known to exist in AGMs^52^, no CD11c clone could be found that is cross reactive with AGMs and sensitive enough for our CyTOF panel. This marker is thus negative in all populations in AGMs. (d) The moderate, wide-spread staining of CD61, which canonically stains platelets, monocytes and macrophages, could be due to platelet debris sticking to other cell types during processing and thus have no biological meaning. However, this marker has also been proposed to be acquired by activated T cells from platelet-derived microvesicles^53^, and in previous studies with cynomolgus macaque blood, we observed subsets of T cells, B cells and NK cells that stain for CD61 (not shown), which could indicate that this is an activation marker in more than just T cells in macaques.

One must also take into consideration necessary technical caveats associated with dosing and cytokine activity before making precise, quantitative conclusions about signaling assays: When available, we used cytokines specific to each species, and we dosed at the same mass concentration (*i.e*. mg/mL) as we dosed for humans as opposed to attempting to coordinate species-specific ECx values. Thus, quantitative comparisons must be verified by close examination of controls and/or by performing additional experiments such as dose-response curves. To that end, we advise comparison with other signaling antibodies, stimulation conditions and/or other cell types within the dataset, which can usually serve as internal controls.

The submitted dataset may be viewed online, and the accompanying article by Fragiadakis & Bjornson, *et al*. provides a detailed analysis of the human data, including correlations with demographics and signaling networks.

## Materials and Methods

### Stimuli

Stimuli used are shown in Table S3. Human and non-primate stimuli were tested in human whole blood over a range of concentrations to select the working concentration. Mouse stimuli were likewise tested in mouse blood. All stimuli were diluted such that the same volume of each would achieve the desired stimulation concentration, then aliquoted into single-use stimuli plates and stored at −80, with the exception of those marked with *, which were dispend at time of use due to storage requirements or the need to adjust the stimulation concentration. All cytokines were tested for endotoxin by the LAL method and verified to contain an amount less than that detectable by our phospho-flow assays (approximately 10 pg/ml) (data not shown).

LPS was commercially prepared by phenol-water extraction and contained small amounts of other bacterial components that activate TLR2. The particular type of LPS (long chain derived from *E. coli* O111:B4) was selected because it is from a pathogenic strain more commonly found in clinical cases^54^.

Rhesus/cynomolgus IL-2 was obtained from the NIH/ NCRR-funded Resource for Nonhuman Primate Immune Reagents.

Gamma-inactivated vegetative *Bacillus anthracis* Ames (ANG-BACI008-VE) was obtained from the Department of Defense Critical Reagents Program through the NIH Biodefense and Emerging Infections Research Resources Repository, NIAID, NIH.

Zaïre Ebolavirus-like particles were used within 48 hours of production, except where noted, and thus different lots were used throughout the course of the project. Particles were produced according to^55^ using calcium phosphate instead of Lipofectamine. Briefly, 1 μg of pCAGGS-GP, 1 μg of pCAGGS-VP35, 1.5 μg of pCAGGS-NP and 1.5 μg of pCAGGS-VP40 (courtesy of R. Johnson *et al.*, NIH NIAID Integrated Research Facility) were transfected into 293T cells by the calcium phosphate method, harvested after 36 hours, then purified through a 20% sucrose cushion at 115,605 x *g* for two hours and stored at +4 degrees. The proteins self-assemble into a structure resembling the native virion^55,56^. These particles are inherently replication-defective and when administered to non-human primates evoke a vaccinating immune response^57–59^, furthermore eliciting type I interferon and proinflammatory cytokine expression^60^. To validate that the production method worked in our hands, preparations were verified by western blot to contain each of the four proteins and by electron microscopy for morphology and quantity. Subsequent preparations were spot-checked by electron microscopy. For particle quantitation and morphological observation, VLP particle preparations were mixed with 110 nm latex spheres (Structure Probe, Inc., West Chester, PA) at a known concentration, absorbed onto copper carbon-Formvar-coated 300 mesh electron microscopy grids prepared in duplicate and stained with uranyl acetate (protocol courtesy of J. Birnbaum, NIH NIAID Integrated Research Facility). Grids were imaged on a JEOL JEM1400 (funded by NIH grant 1Z10RR02678001). Due to the short shelf-life of the particles, grids were generally imaged after use as a stimulus.

### Blood

Venous human blood was obtained from the Stanford Blood Center, AllCells Inc. (Alameda, CA) (exempt, non-human subjects research) or from volunteers from the Stanford community under an IRB-approved protocol (#28289).

Macaque (*Macaca mulatta* and *M. fascicularis*) blood was obtained from Valley Biosystems from conscious (not sedated), captive-born, Chinese-origin animals. African green monkey (*Cercopithecus aethiops*) blood was obtained from Worldwide Primates and Bioreclamation, LLC. The African green monkeys were wild-born in St. Kitts; age was estimated by capture date (assuming two years old at time of capture). Mice were obtained from Charles River Laboratories. Animal work was done under an approved animal care and usage committee protocol (#26675).

Health reports for the macaques were obtained and are available in the experiment data repository.

Complete blood counts (CBCs) were performed on a Sysmex XT-2000iv with the veterinary software module.

### Antibodies

Purified antibodies were purchased and conjugated in-house using DVS/Fluidigim MaxPar X8 metal conjugation kits (Tables S4 – S7). All antibodies were titrated for optimal signal-to-noise ratio, then re-confirmed in at least two different individuals per species (three humans, two cynomolgus macaques, two rhesus macaques, three mice). All conjugations and titrations were well-documented, and records are available in the experiment data repository. Finally, antibodies were lyophilized into LyoSpheres by BioLyph LLC (Hopkins, MN) with excipient B144 as 4x cocktails. CyTOF antibody LyoSpheres were stress-tested for over one year and found to have no significant change in staining (not shown).

### Stimulation and Staining

Stimulation and staining was carried out on a custom automation platform consisting of an Agilent Bravo pipetting robot, Agilent BenchBot robotic arm, Peak KiNeDx robotic arm, Thermo Cytomat C2 incubator, BioTek ELx405-UVSD aspirator/dispenser, BioTek MultiFlo FX four-reagent dispenser, Q.Instruments microplate shakers, Velocity11 VSpin centrifuges and a custom chilling system contained in a negative-pressure biosafety enclosure. The VWorks robotic programs and logs from protocol runs are available upon request.

Whole blood was stimulated by mixing with stimuli and incubating in a humidified 37-degree, 5% CO_2_ incubator for 15 minutes. Blood was fixed for 10 minutes at room temperature with 1.6% paraformaldehyde (PFA, Electron Microscopy Sciences) and lysed with 0.1% Triton-X100 in PBS for 30 minutes at room temperature per^61^. Cells were washed twice with PBS, then each donor’s 16 conditions were barcoded according to^62,63^. Briefly, cells were permeabilized with 0.02% saponin, then stained with unique combinations of functionalized, stable palladium isotopes. The stimulation plate containing 6 donors x 16 conditions was then reduced to 6 wells, each containing the 16 conditions for one donor. Cells were washed once with staining media (CSM: 0.2% BSA in PBS with 0.02% sodium azide), blocked with human (humans, NHPs) or mouse (mice) TruStain FcX block (Biolegend) for 10 minutes at room temperature with shaking, then stained with rehydrated extracellular LyoSpheres for 30 minutes at room temperature with shaking in a final volume of 240 μl. (See Table S4 - S7 for final staining concentrations.) Cells were washed once, then permeabilized in >90% methanol at 4 degrees C for 20 minutes. Cells were washed four times, then stained with intracellular lyospheres for 60 minutes at room temperature with shaking. Cells were washed once, then placed into 1.6% PFA and 0.1 μM natural iridium intercalator (Fluidigm) in PBS at 4 degrees C until acquisition on a CyTOF. With few exceptions, cells were acquired within seven days of staining. From prior validation experiments, this amount of time imparts no significant effect on staining.

### Acquisition

Prior to running, cells were washed twice with water. Samples were acquired on a single DVS/Fluidigm CyTOF 2 fitted with a Super Sampler sample introduction system (Victorian Airship & Scientific Apparatus LLC). QC reports were run on the CyTOF between every barcoded sample. Prior to beginning acquisition, the instrument must have demonstrated Tb159 dual counts > 1,000,000 and oxidation < 3%; if the instrument failed those criteria, it was cleaned, tuned or repaired as necessary. Approximately 4,800,000 events were acquired per sample.

## Disclaimer and Acknowledgements

The research discussed in this article was supported in part by the U.S. Food and Drug Administration (Contract No. HHSF223201210194C). Additional support was provided by NIH awards 5R01CA18496804, 5R25CA18099304, 1R01GM10983604, 5UH2AR06767603, 1R01NS08953304 and R01HL120724, and FDA contract HHSF223201610018C. Z.B.B.H. was supported in part by NIH grant T32GM007276. G.K.F. was supported in part by a Stanford Bio-X graduate research fellowship and NIH grant T32GM007276. M.H.S. was supported in part by NIH grant DP5OD023056. This article reflects the views of the authors and should not be construed to represent the U.S. Food and Drug Administration or NIH’s views or policies.

### Author contributions

Z.B.B.H. generated the data, performed the analysis and wrote the manuscript. G.K.F. contributed to data generation and edited the manuscript. M.H.S. contributed to data generation and edited the manuscript. D.Madhireddy contributed to data generation and reagent optimization. D.McIlwain edited the manuscript and advised analysis. G.P.N. advised the study and edited the manuscript.

### Declaration of interests

Z.B.B.H. is involved in the commercial development of the platform used to host the data presented here, although the data presented here will always be freely accessible.

