## Supplemental Figures and Tables for "A comprehensive atlas of immunological differences between humans, mice and non-human primates"

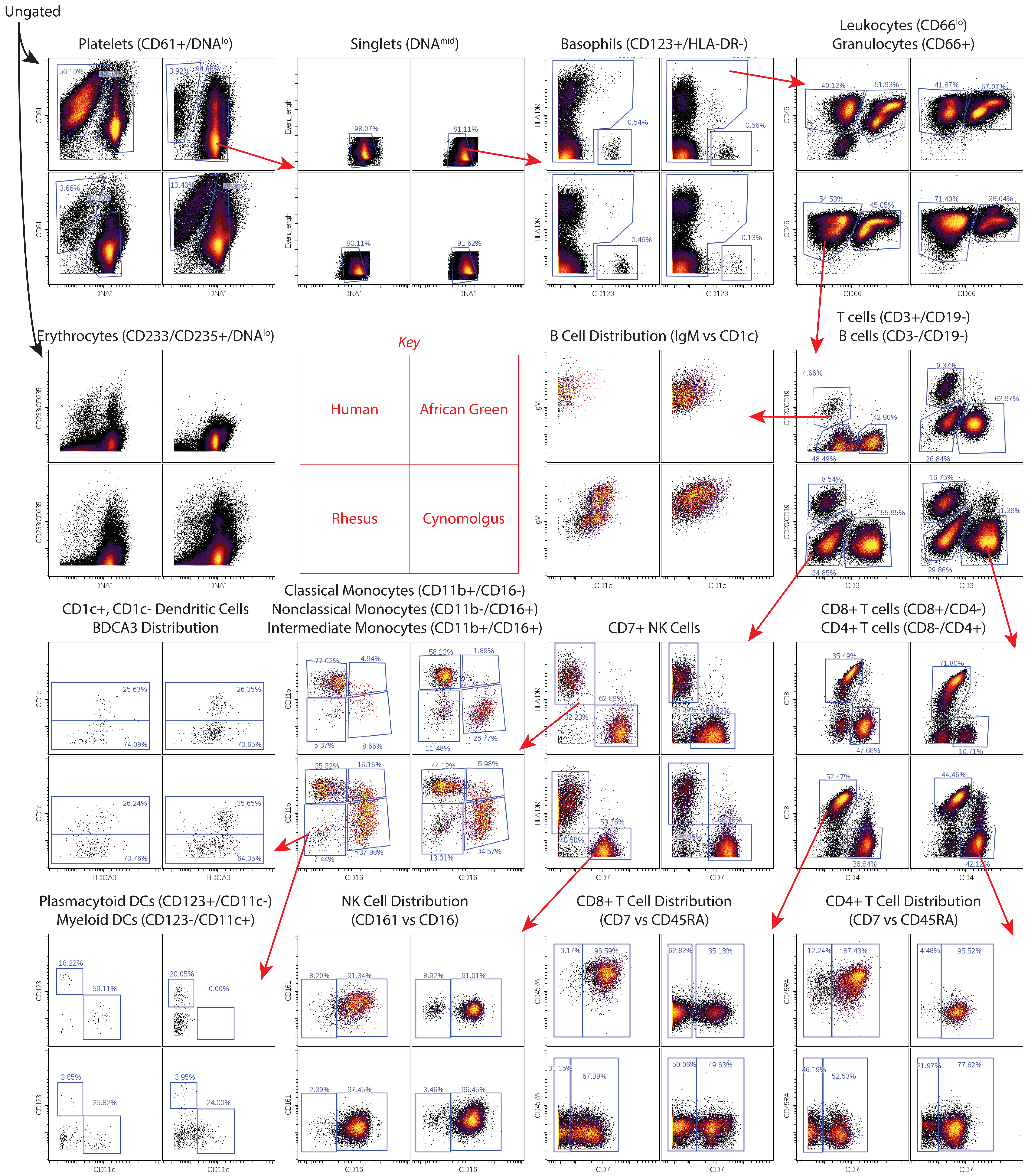
Ungated
Platelets (CD61+/DNA^lo)
Singlets (DNA^mid)
Basophils (CD123+/HLA-DR-)
Leukocytes (CD66^lo)
Granulocytes (CD66+)
Erythrocytes (CD233/CD235+/DNA^lo)
Key
Human
African Green
Rhesus
Cynomolgus
B Cell Distribution (IgM vs CD1c)
T cells (CD3+/CD19-)
B cells (CD3-/CD19-)
CD1c+, CD1c- Dendritic Cells
BDCA3 Distribution
Classical Monocytes (CD11b+/CD16-)
Nonclassical Monocytes (CD11b-/CD16+)
Intermediate Monocytes (CD11b+/CD16+)
CD7+ NK Cells
CD8+ T cells (CD8+/CD4-)
CD4+ T cells (CD8-/CD4+)
Plasmacytoid DCs (CD123+/CD11c-)
Myeloid DCs (CD123-/CD11c+)
NK Cell Distribution
(CD161 vs CD16)
CD8+ T Cell Distribution
(CD7 vs CD45RA)
CD4+ T Cell Distribution
(CD7 vs CD45RA)
CD61
DNA1
Event_length
HLA-DR
CD123
CD45
CD66
CD233/CD235
IgM
CD1c
CD20/CD19
CD3
CD11b
CD16
CD7
CD8
CD4
BDCA3
CD161
CD45RA
CD11c

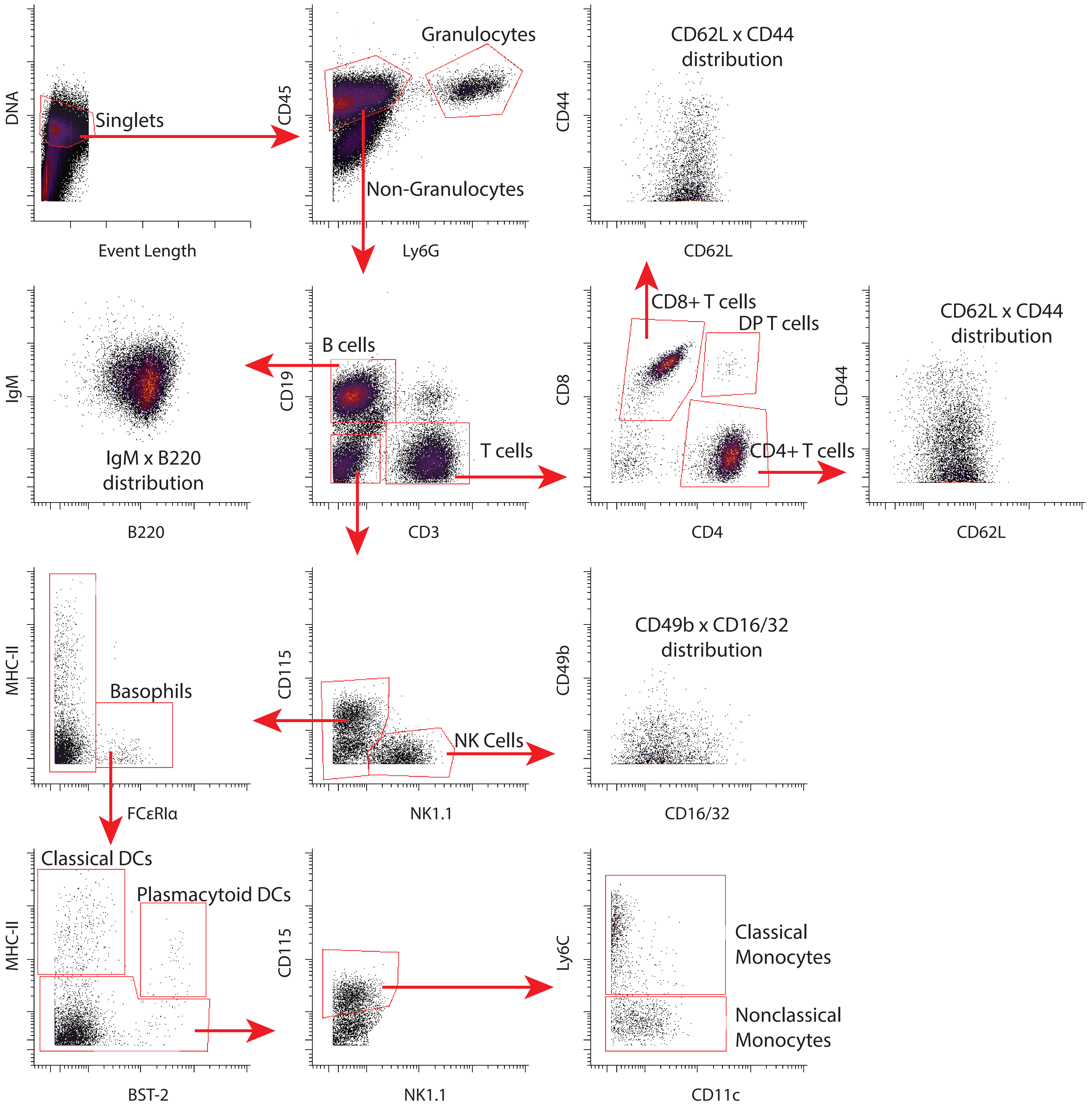

DNA
Singlets
Event Length
CD45
Granulocytes
Non-Granulocytes
Ly6G
CD44
CD62L x CD44
distribution
CD62L
IgM
IgM x B220
distribution
B220
CD19
B cells
T cells
CD3
CD8
CD8+ T cells
DP T cells
CD4+ T cells
CD4
CD44
CD62L x CD44
distribution
CD62L
MHC-II
Basophils
FCεRIα
CD115
NK Cells
NK1.1
CD49b
CD49b x CD16/32
distribution
CD16/32
MHC-II
Classical DCs
Plasmacytoid DCs
BST-2
CD115
NK1.1
Ly6C
Classical
Monocytes
Nonclassical
Monocytes
CD11c

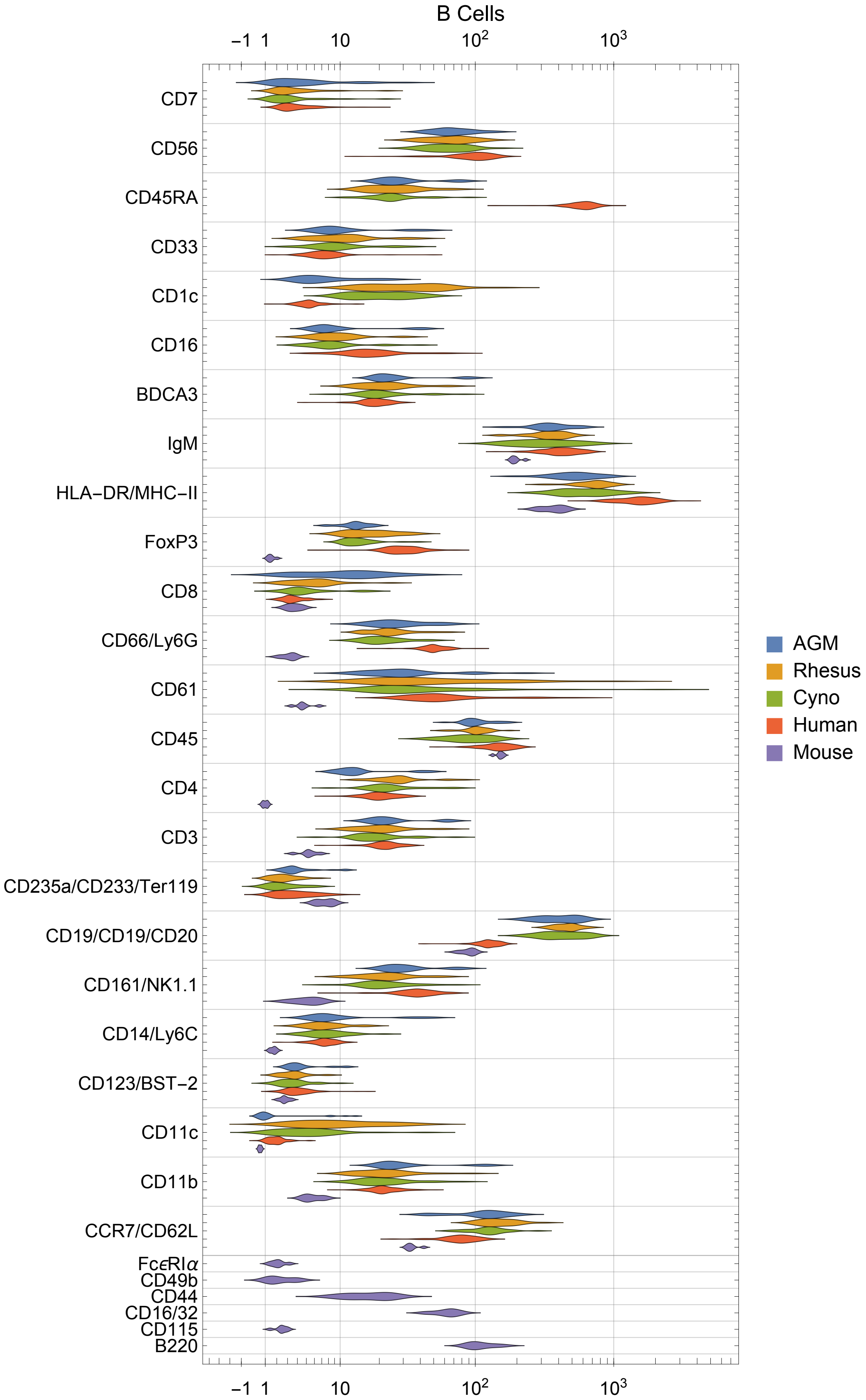
B Cells
−1 1 10 10^2 10^3
CD7
CD56
CD45RA
CD33
CD1c
CD16
BDCA3
IgM
HLA−DR/MHC−II
FoxP3
CD8
CD66/Ly6G
CD61
CD45
CD4
CD3
CD235a/CD233/Ter119
CD19/CD19/CD20
CD161/NK1.1
CD14/Ly6C
CD123/BST−2
CD11c
CD11b
CCR7/CD62L
FcεRIα
CD49b
CD44
CD16/32
CD115
B220
AGM
Rhesus
Cyno
Human
Mouse

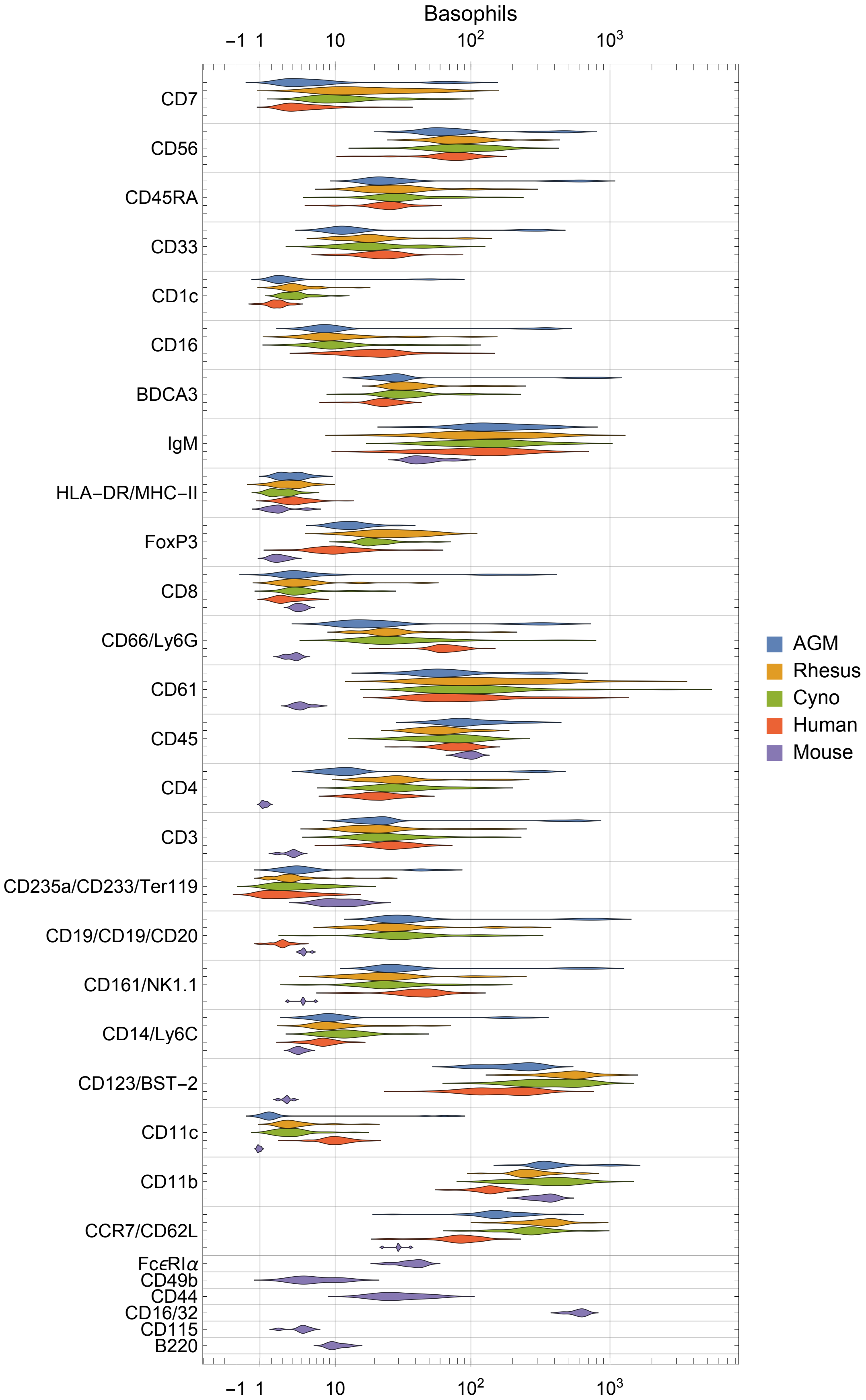

Basophils
−1 1 10 10^2 10^3
CD7
CD56
CD45RA
CD33
CD1c
CD16
BDCA3
IgM
HLA−DR/MHC−II
FoxP3
CD8
CD66/Ly6G
CD61
CD45
CD4
CD3
CD235a/CD233/Ter119
CD19/CD19/CD20
CD161/NK1.1
CD14/Ly6C
CD123/BST−2
CD11c
CD11b
CCR7/CD62L
FcεRIα
CD49b
CD44
CD16/32
CD115
B220
AGM
Rhesus
Cyno
Human
Mouse

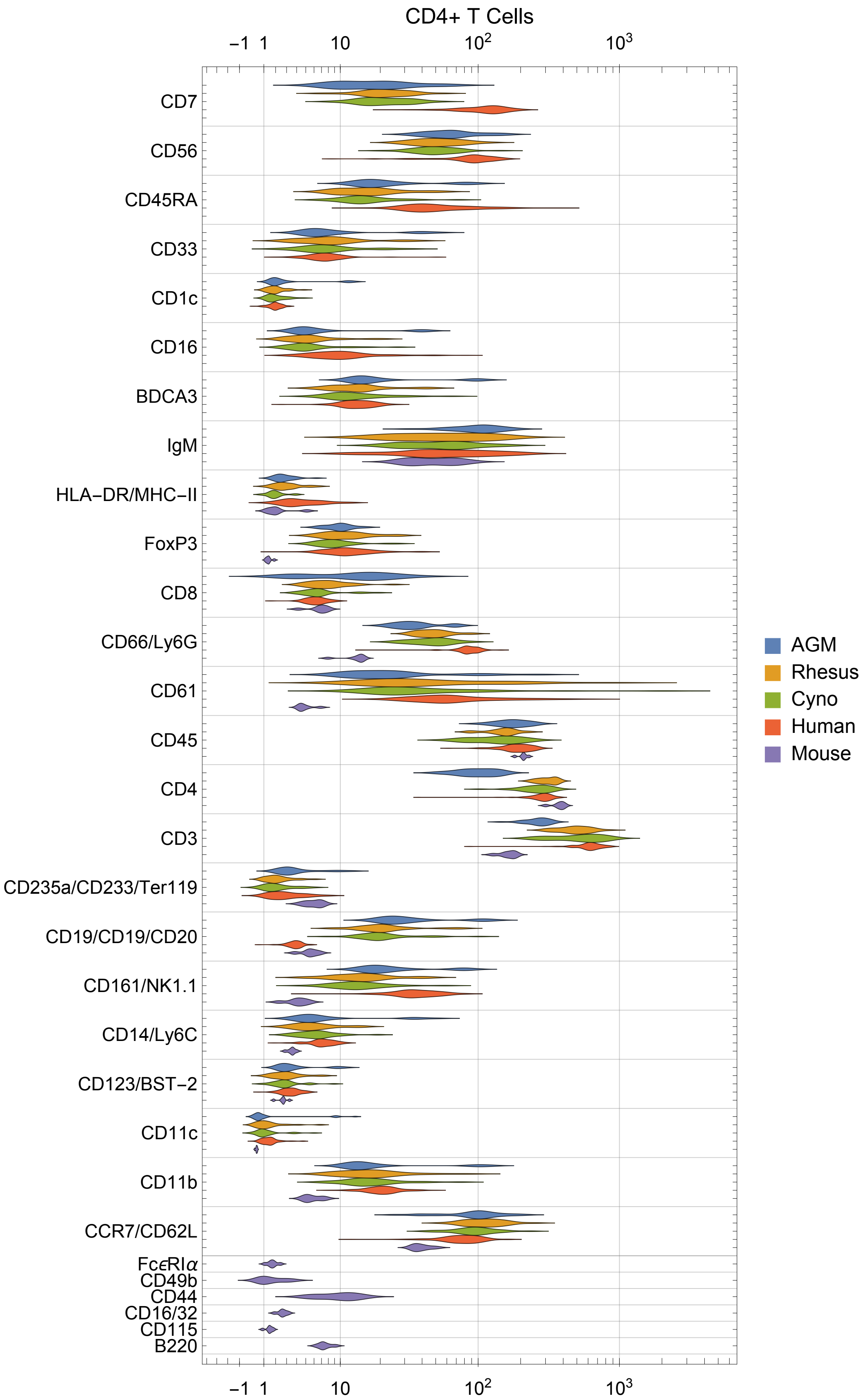

CD4+ T Cells
−1
1
10
$10^2$
$10^3$
CD7
CD56
CD45RA
CD33
CD1c
CD16
BDCA3
IgM
HLA−DR/MHC−II
FoxP3
CD8
CD66/Ly6G
CD61
CD45
CD4
CD3
CD235a/CD233/Ter119
CD19/CD19/CD20
CD161/NK1.1
CD14/Ly6C
CD123/BST−2
CD11c
CD11b
CCR7/CD62L
FcεRIα
CD49b
CD44
CD16/32
CD115
B220
AGM
Rhesus
Cyno
Human
Mouse

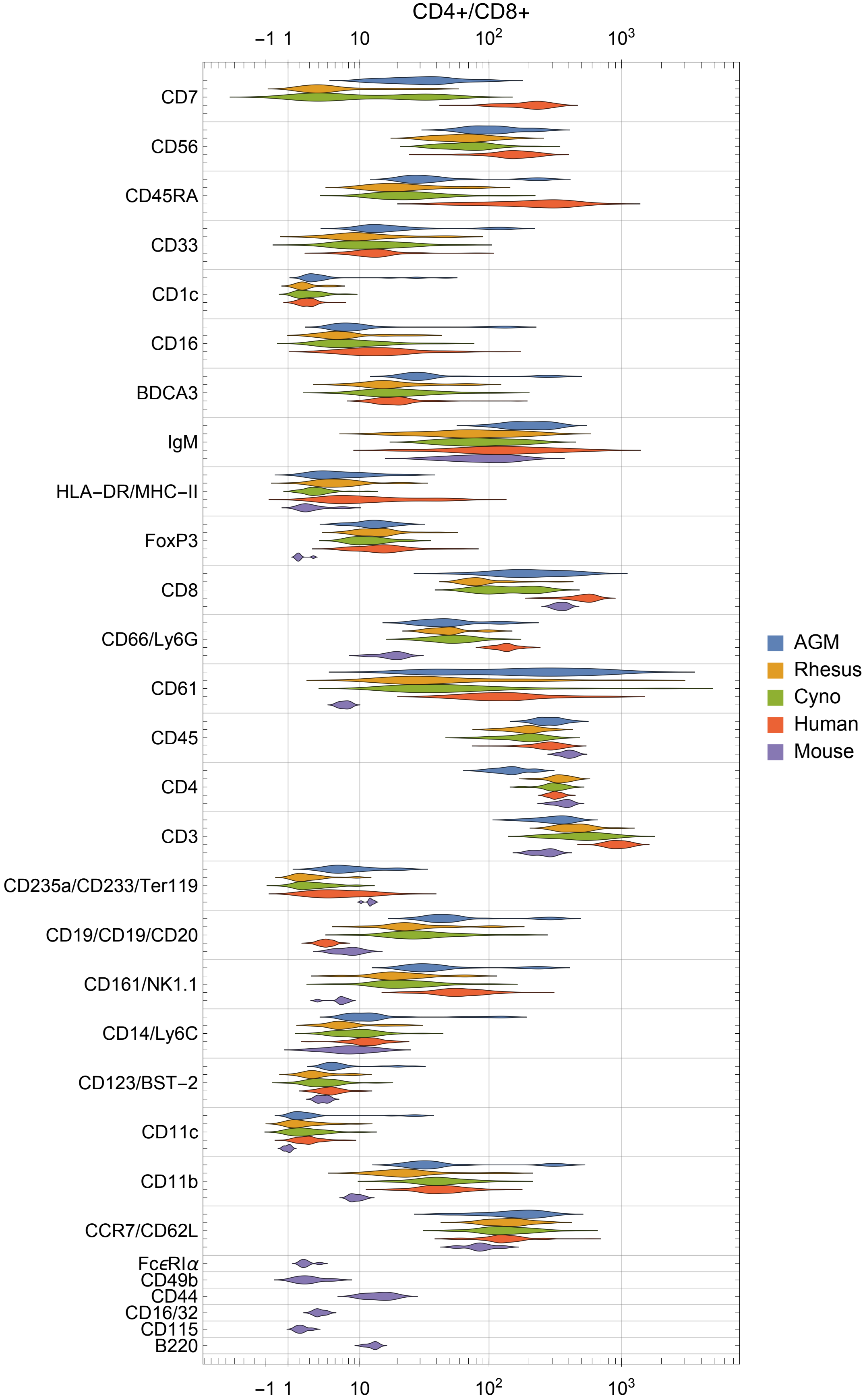

CD4+/CD8+
−1 1
10
$10^2$
$10^3$
CD7
CD56
CD45RA
CD33
CD1c
CD16
BDCA3
IgM
HLA−DR/MHC−II
FoxP3
CD8
CD66/Ly6G
CD61
CD45
CD4
CD3
CD235a/CD233/Ter119
CD19/CD19/CD20
CD161/NK1.1
CD14/Ly6C
CD123/BST−2
CD11c
CD11b
CCR7/CD62L
FcϵRIα
CD49b
CD44
CD16/32
CD115
B220
AGM
Rhesus
Cyno
Human
Mouse

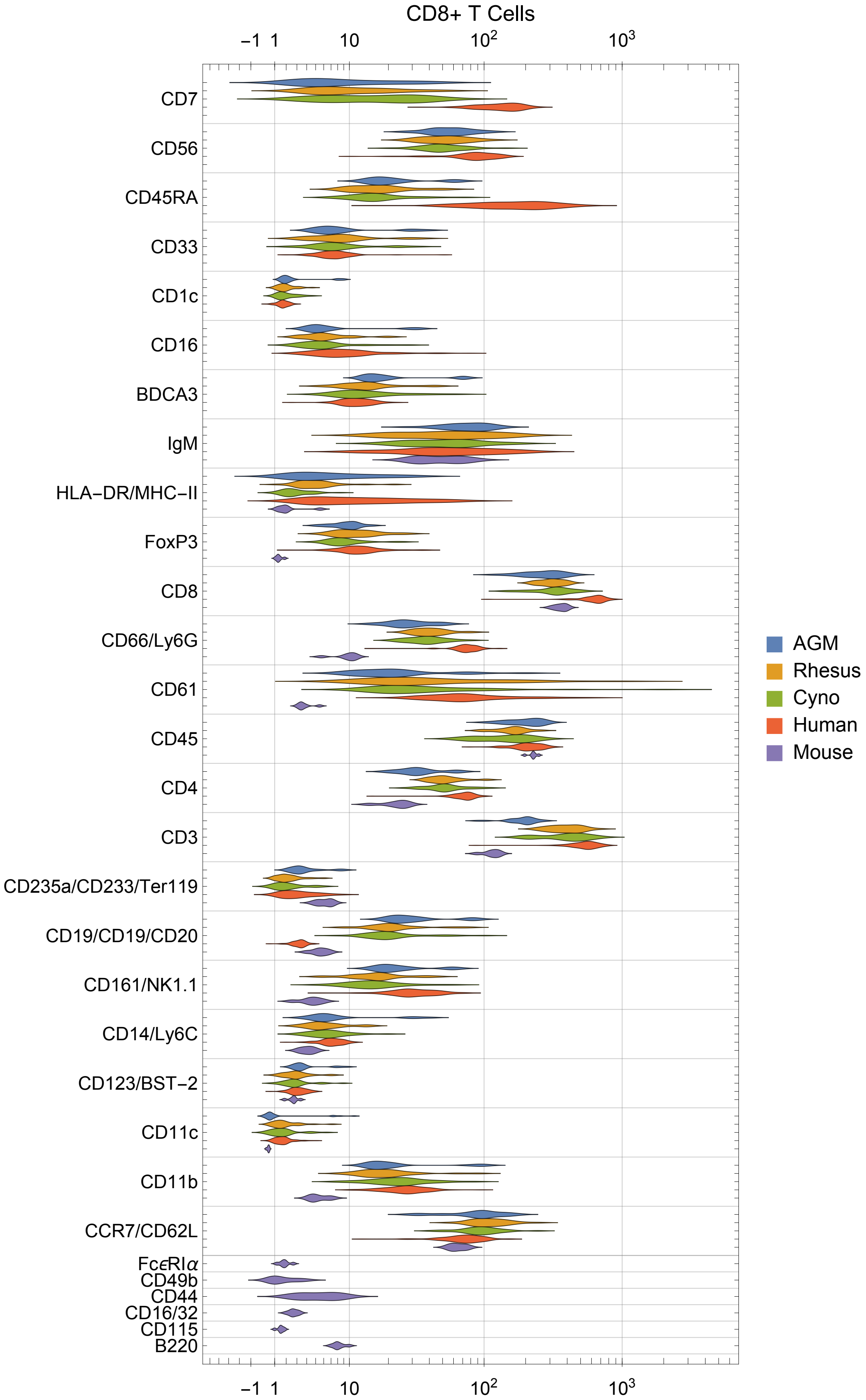
CD8+ T Cells
−1
1
10
$10^2$
$10^3$
CD7
CD56
CD45RA
CD33
CD1c
CD16
BDCA3
IgM
HLA−DR/MHC−II
FoxP3
CD8
CD66/Ly6G
CD61
CD45
CD4
CD3
CD235a/CD233/Ter119
CD19/CD19/CD20
CD161/NK1.1
CD14/Ly6C
CD123/BST−2
CD11c
CD11b
CCR7/CD62L
FcϵRIα
CD49b
CD44
CD16/32
CD115
B220
AGM
Rhesus
Cyno
Human
Mouse

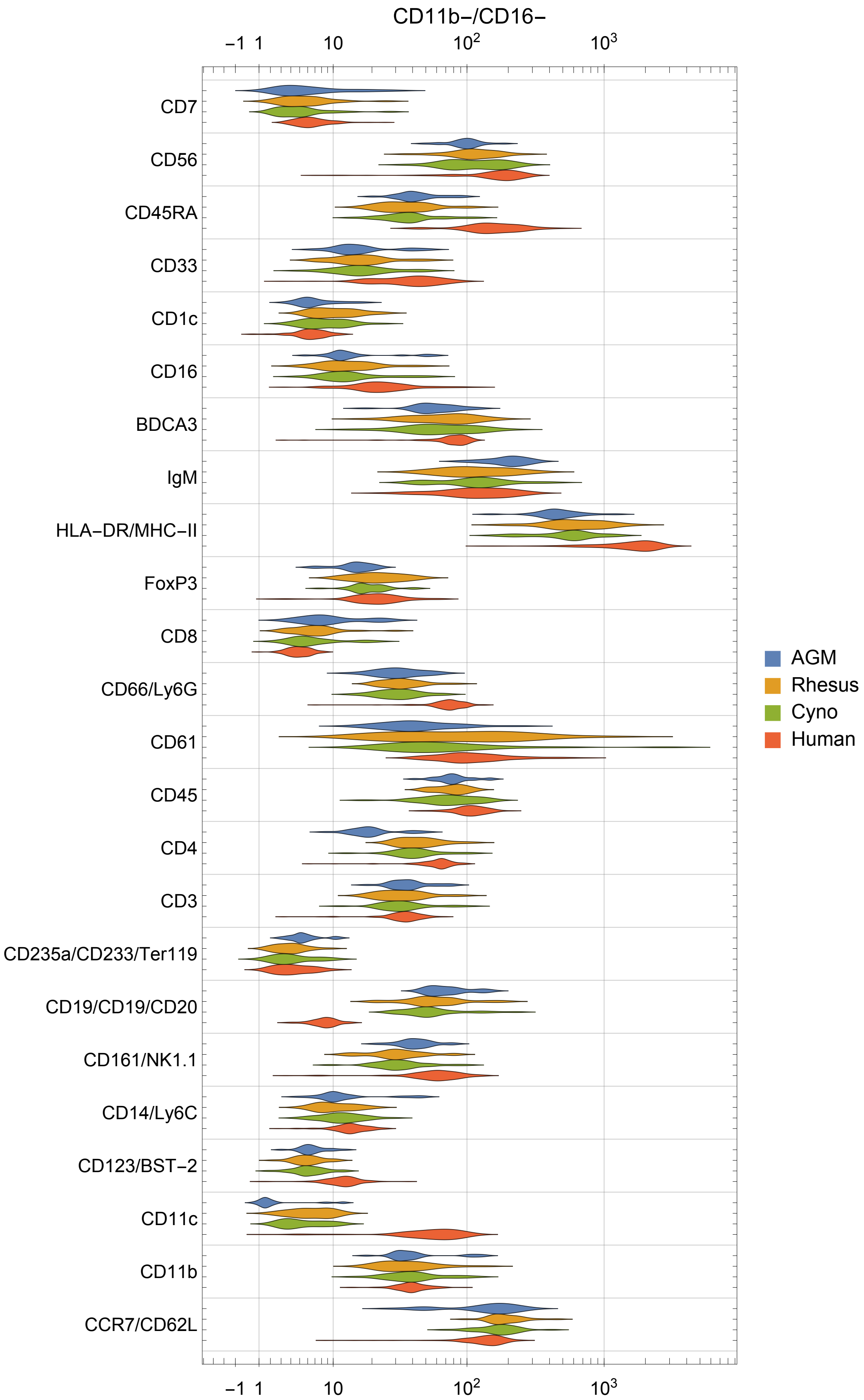

CD11b−/CD16−
−1 1 10 10^2 10^3
CD7
CD56
CD45RA
CD33
CD1c
CD16
BDCA3
IgM
HLA−DR/MHC−II
FoxP3
CD8
CD66/Ly6G
CD61
CD45
CD4
CD3
CD235a/CD233/Ter119
CD19/CD19/CD20
CD161/NK1.1
CD14/Ly6C
CD123/BST−2
CD11c
CD11b
CCR7/CD62L
AGM
Rhesus
Cyno
Human

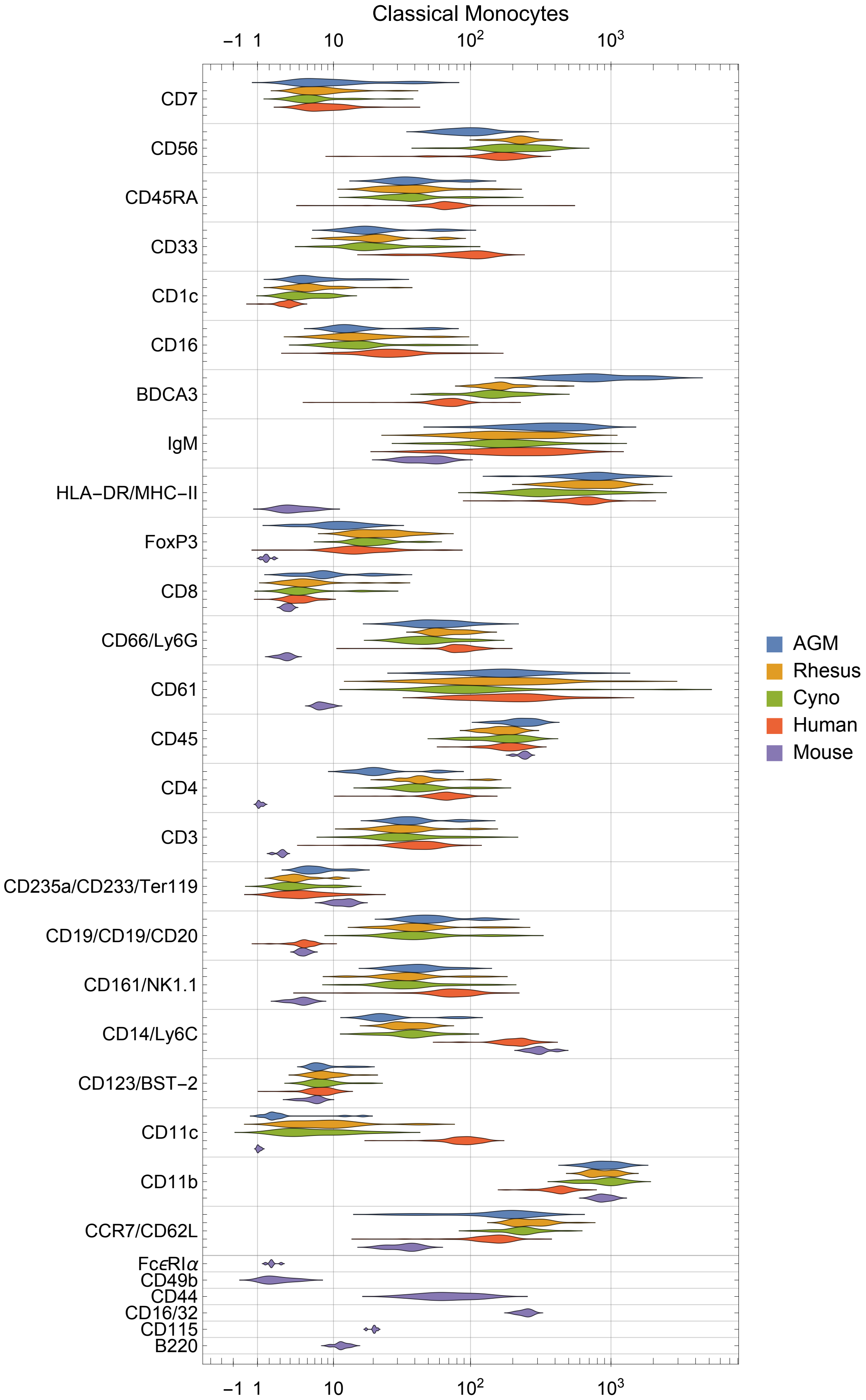

Classical Monocytes
-1
1
10
$10^2$
$10^3$
CD7
CD56
CD45RA
CD33
CD1c
CD16
BDCA3
IgM
HLA-DR/MHC-II
FoxP3
CD8
CD66/Ly6G
CD61
CD45
CD4
CD3
CD235a/CD233/Ter119
CD19/CD19/CD20
CD161/NK1.1
CD14/Ly6C
CD123/BST-2
CD11c
CD11b
CCR7/CD62L
FcϵRIα
CD49b
CD44
CD16/32
CD115
B220
AGM
Rhesus
Cyno
Human
Mouse

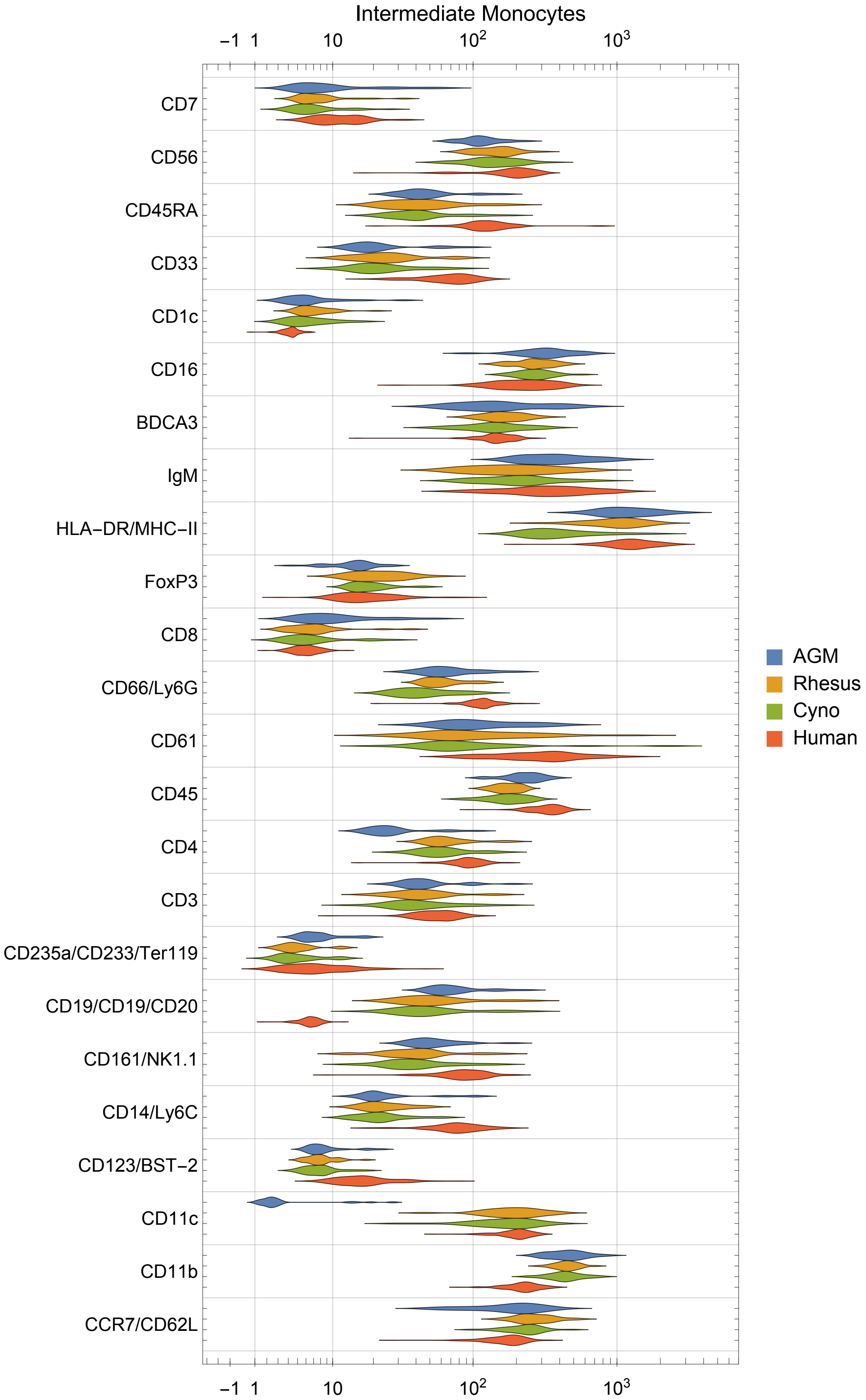

Intermediate Monocytes
−1
1
10
$10^2$
$10^3$
CD7
CD56
CD45RA
CD33
CD1c
CD16
BDCA3
IgM
HLA−DR/MHC−II
FoxP3
CD8
CD66/Ly6G
CD61
CD45
CD4
CD3
CD235a/CD233/Ter119
CD19/CD19/CD20
CD161/NK1.1
CD14/Ly6C
CD123/BST−2
CD11c
CD11b
CCR7/CD62L
AGM
Rhesus
Cyno
Human

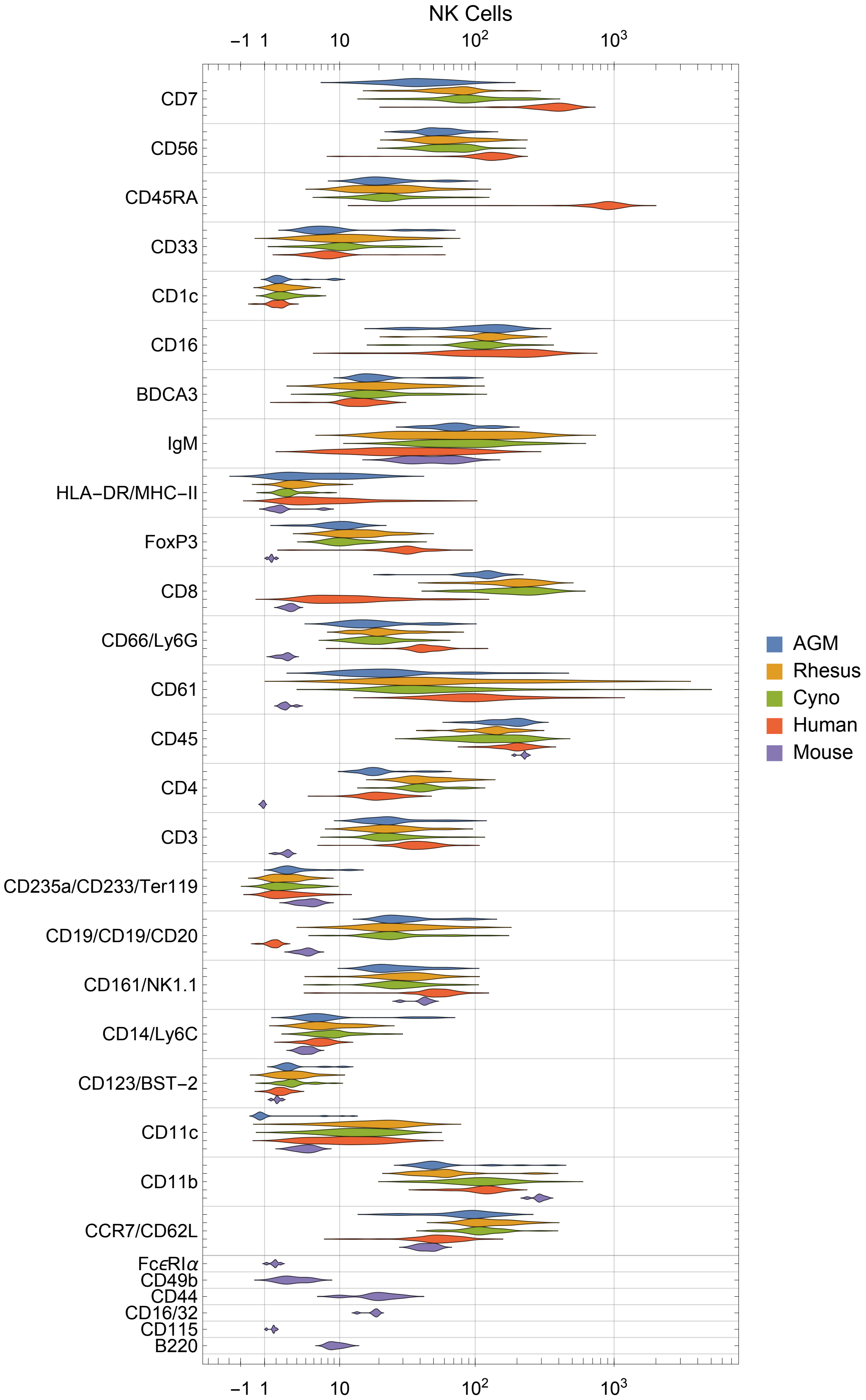

NK Cells
-1
1
10
10^2
10^3
CD7
CD56
CD45RA
CD33
CD1c
CD16
BDCA3
IgM
HLA-DR/MHC-II
FoxP3
CD8
CD66/Ly6G
CD61
CD45
CD4
CD3
CD235a/CD233/Ter119
CD19/CD19/CD20
CD161/NK1.1
CD14/Ly6C
CD123/BST-2
CD11c
CD11b
CCR7/CD62L
FcεRIα
CD49b
CD44
CD16/32
CD115
B220
AGM
Rhesus
Cyno
Human
Mouse

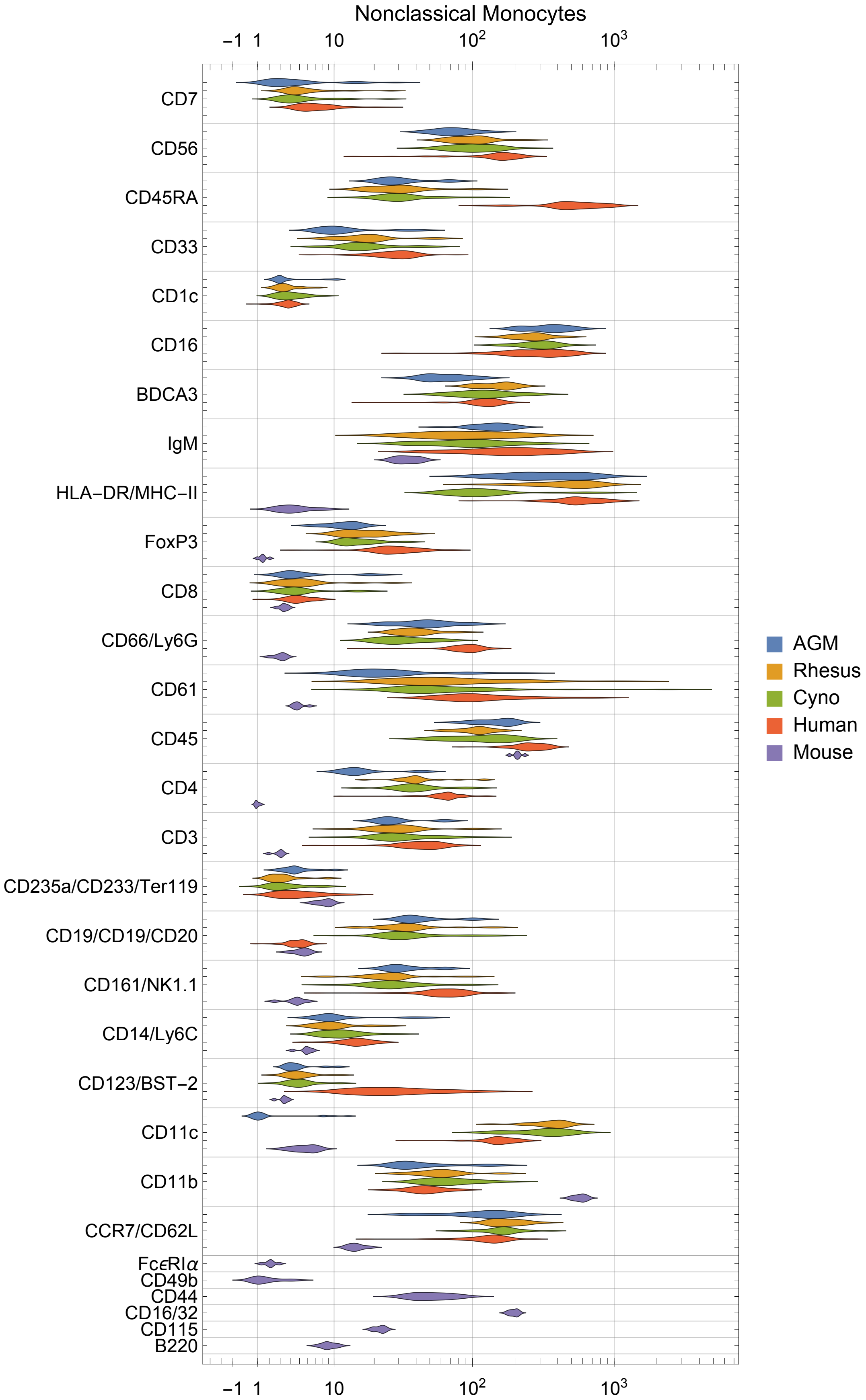

Nonclassical Monocytes
−1
1
10
$10^2$
$10^3$
CD7
CD56
CD45RA
CD33
CD1c
CD16
BDCA3
IgM
HLA−DR/MHC−II
FoxP3
CD8
CD66/Ly6G
CD61
CD45
CD4
CD3
CD235a/CD233/Ter119
CD19/CD19/CD20
CD161/NK1.1
CD14/Ly6C
CD123/BST−2
CD11c
CD11b
CCR7/CD62L
FcϵRIα
CD49b
CD44
CD16/32
CD115
B220
AGM
Rhesus
Cyno
Human
Mouse

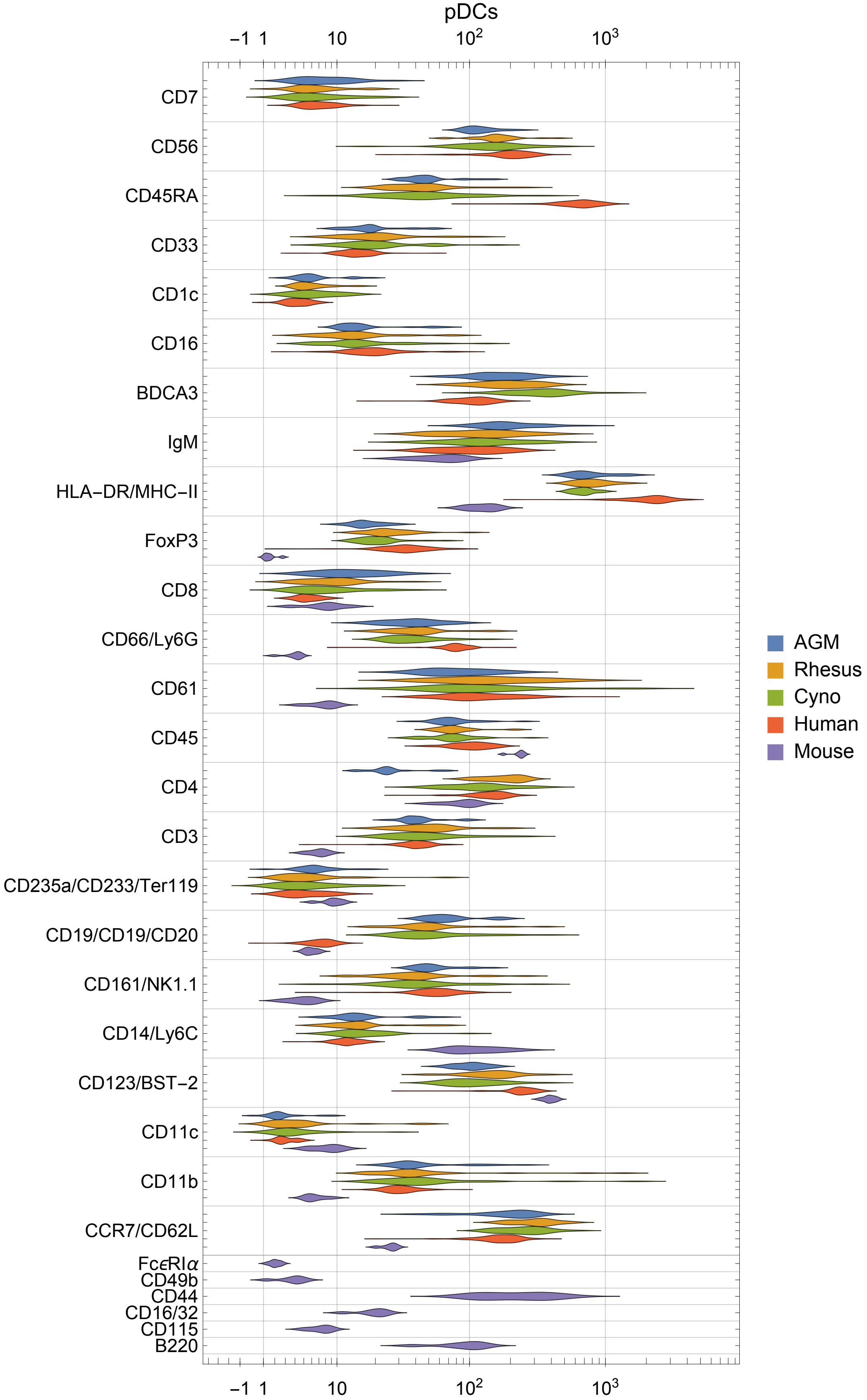
pDCs
-1
1
10
$10^2$
$10^3$
CD7
CD56
CD45RA
CD33
CD1c
CD16
BDCA3
IgM
HLA-DR/MHC-II
FoxP3
CD8
CD66/Ly6G
CD61
CD45
CD4
CD3
CD235a/CD233/Ter119
CD19/CD19/CD20
CD161/NK1.1
CD14/Ly6C
CD123/BST-2
CD11c
CD11b
CCR7/CD62L
FcεRIα
CD49b
CD44
CD16/32
CD115
B220
AGM
Rhesus
Cyno
Human
Mouse

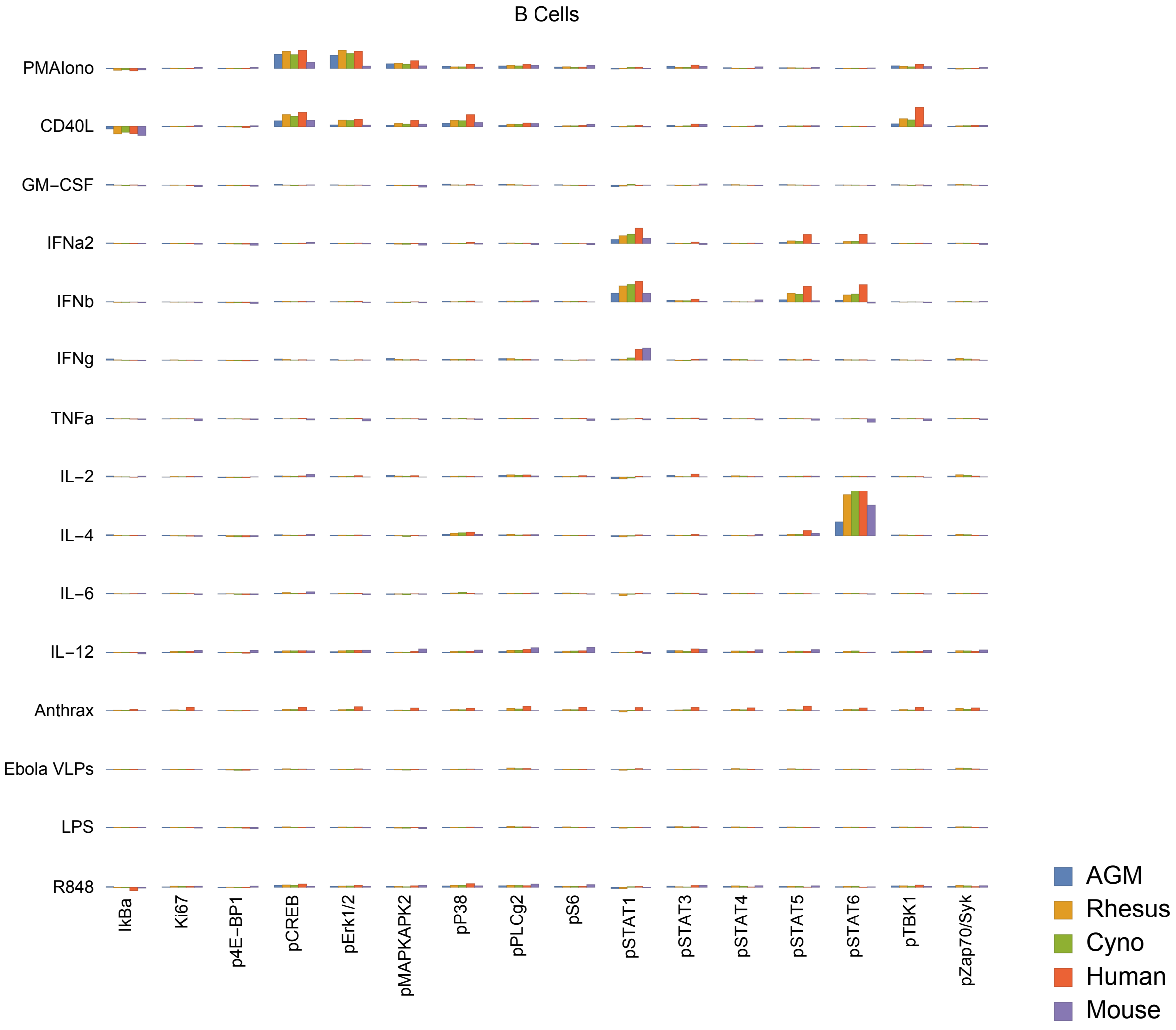
B Cells
PMAIono
CD40L
GM−CSF
IFNa2
IFNb
IFNg
TNFa
IL−2
IL−4
IL−6
IL−12
Anthrax
Ebola VLPs
LPS
R848
IkBa
Ki67
p4E−BP1
pCREB
pErk1/2
pMAPKAPK2
pP38
pPLCg2
pS6
pSTAT1
pSTAT3
pSTAT4
pSTAT5
pSTAT6
pTBK1
pZap70/Syk
AGM
Rhesus
Cyno
Human
Mouse

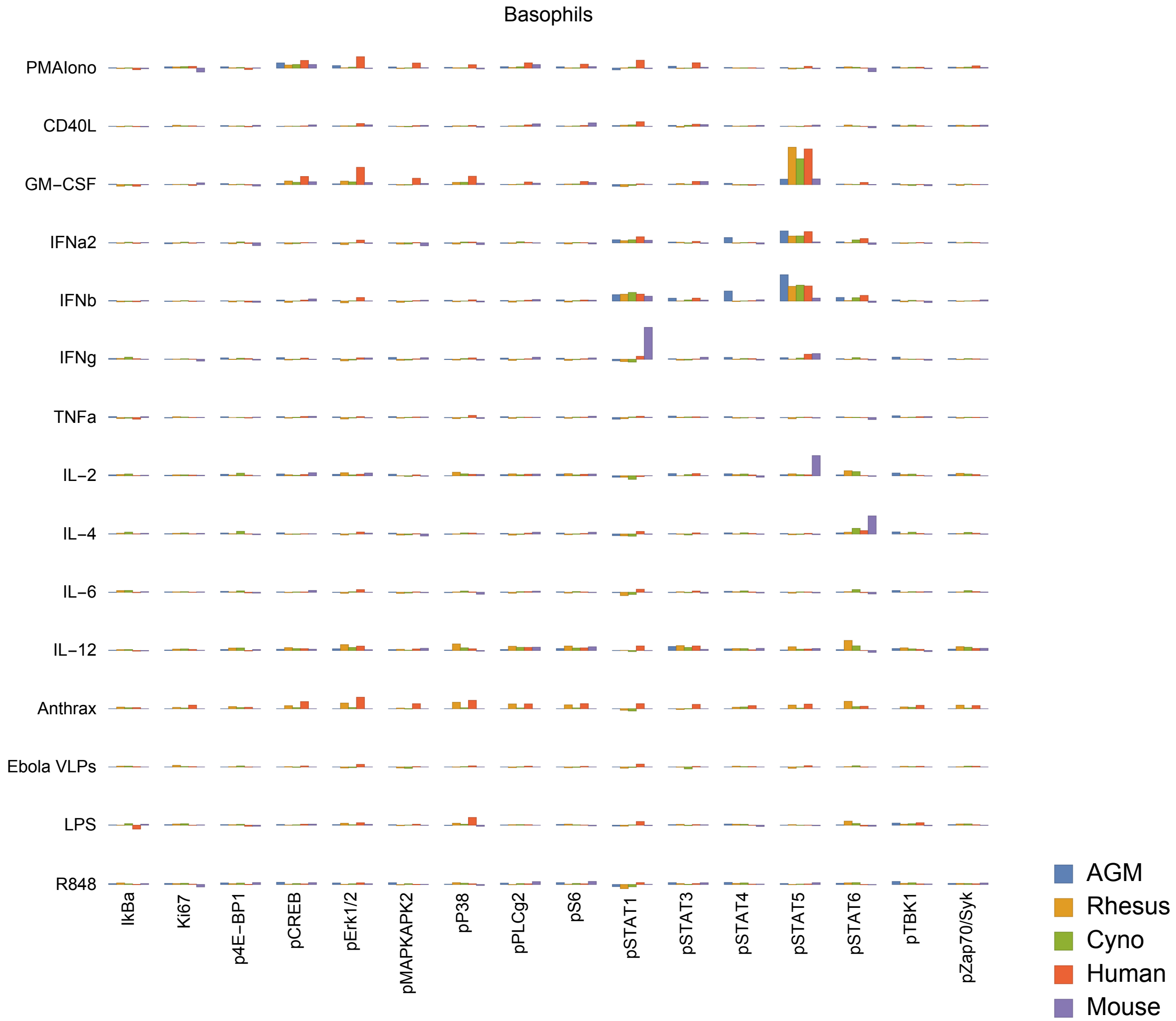

Basophils
PMAIono
CD40L
GM−CSF
IFNa2
IFNb
IFNg
TNFa
IL−2
IL−4
IL−6
IL−12
Anthrax
Ebola VLPs
LPS
R848
IkBa
Ki67
p4E−BP1
pCREB
pErk1/2
pMAPKAPK2
pP38
pPLCg2
pS6
pSTAT1
pSTAT3
pSTAT4
pSTAT5
pSTAT6
pTBK1
pZap70/Syk
AGM
Rhesus
Cyno
Human
Mouse

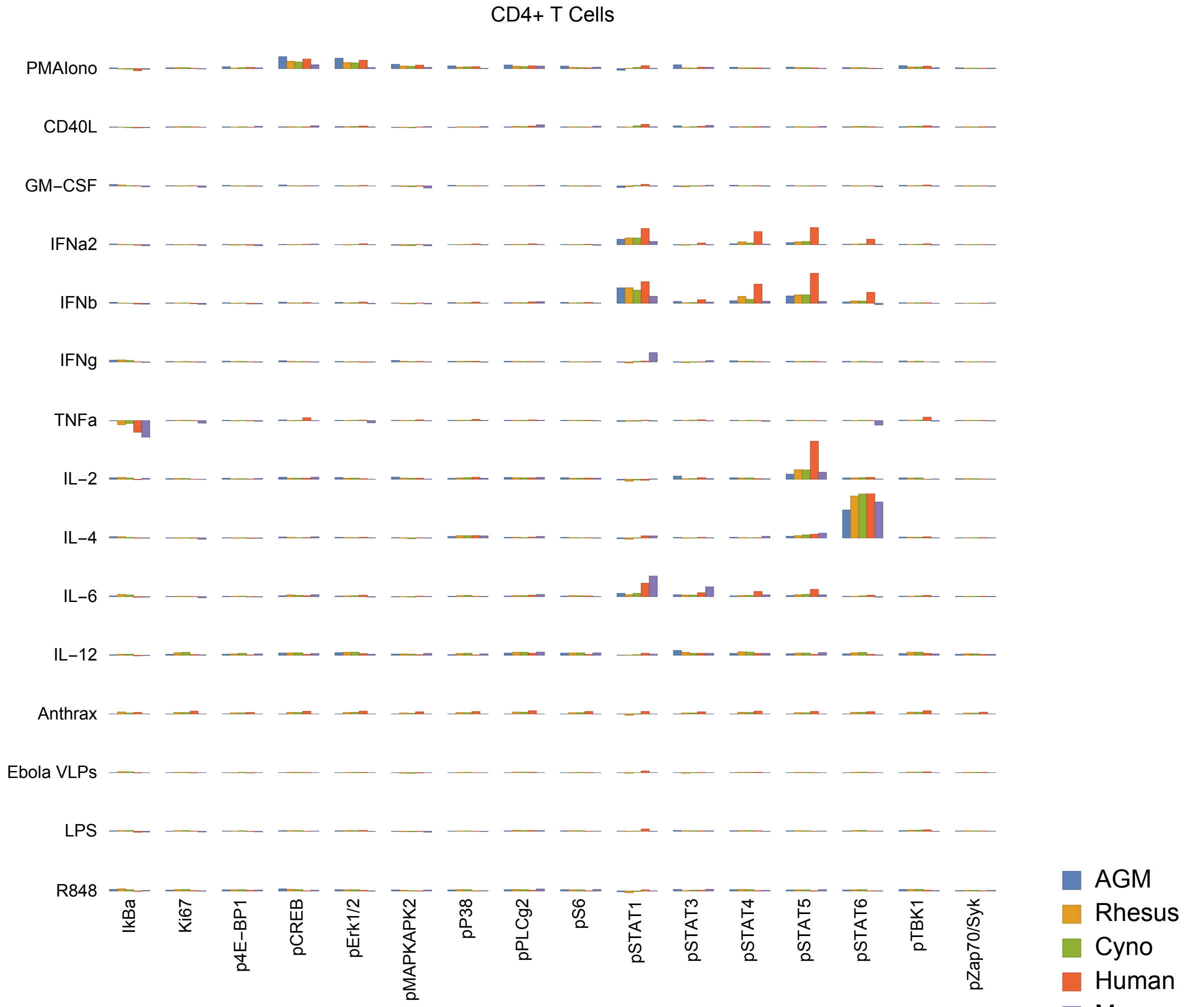
CD4+ T Cells
PMAIono
CD40L
GM−CSF
IFNa2
IFNb
IFNg
TNFa
IL−2
IL−4
IL−6
IL−12
Anthrax
Ebola VLPs
LPS
R848
IkBa
Ki67
p4E−BP1
pCREB
pErk1/2
pMAPKAPK2
pP38
pPLCg2
pS6
pSTAT1
pSTAT3
pSTAT4
pSTAT5
pSTAT6
pTBK1
pZap70/Syk
AGM
Rhesus
Cyno
Human
Mouse

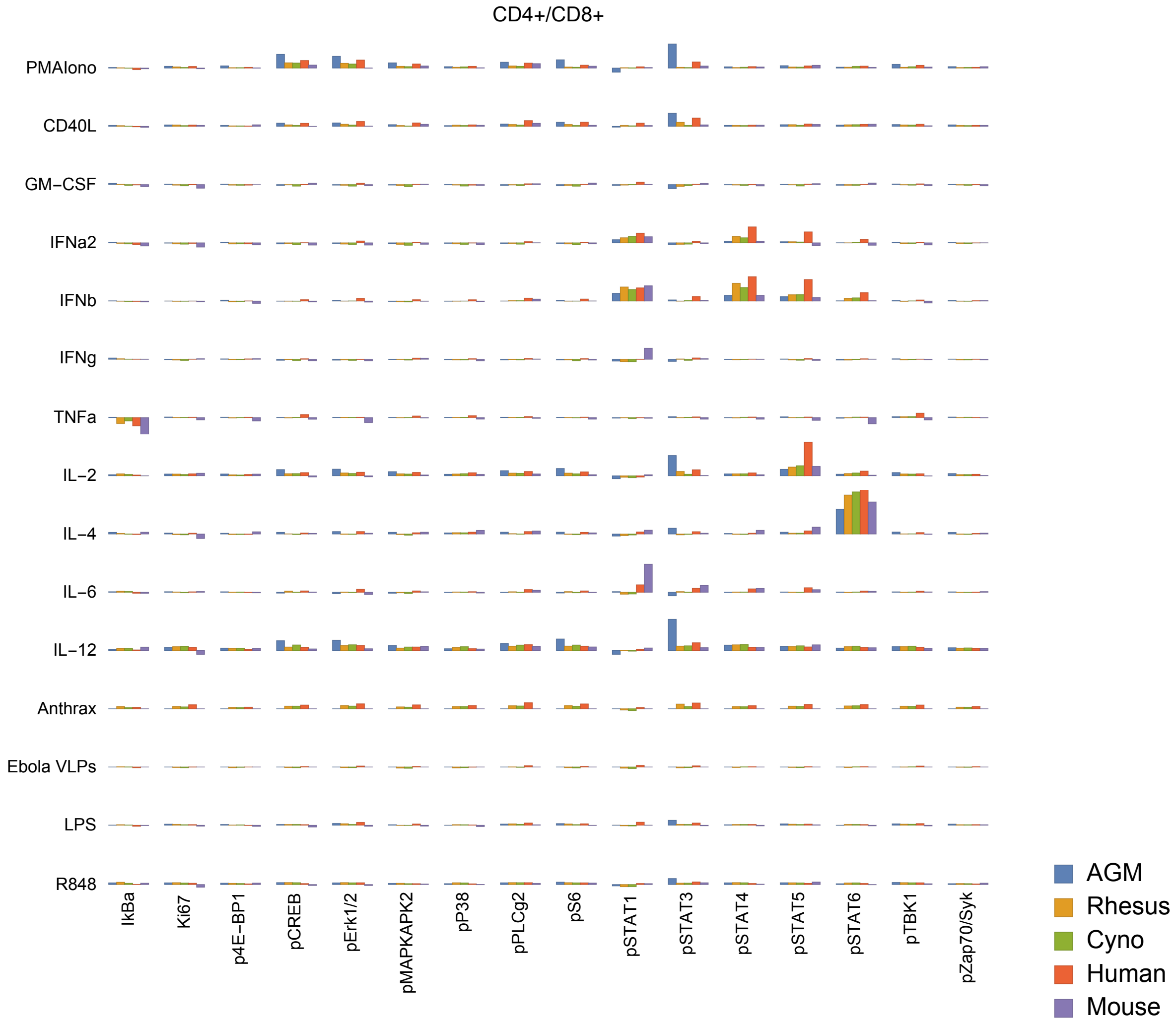
CD4+/CD8+
PMAIono
CD40L
GM−CSF
IFNa2
IFNb
IFNg
TNFa
IL−2
IL−4
IL−6
IL−12
Anthrax
Ebola VLPs
LPS
R848
IkBa
Ki67
p4E−BP1
pCREB
pErk1/2
pMAPKAPK2
pP38
pPLCg2
pS6
pSTAT1
pSTAT3
pSTAT4
pSTAT5
pSTAT6
pTBK1
pZap70/Syk
AGM
Rhesus
Cyno
Human
Mouse

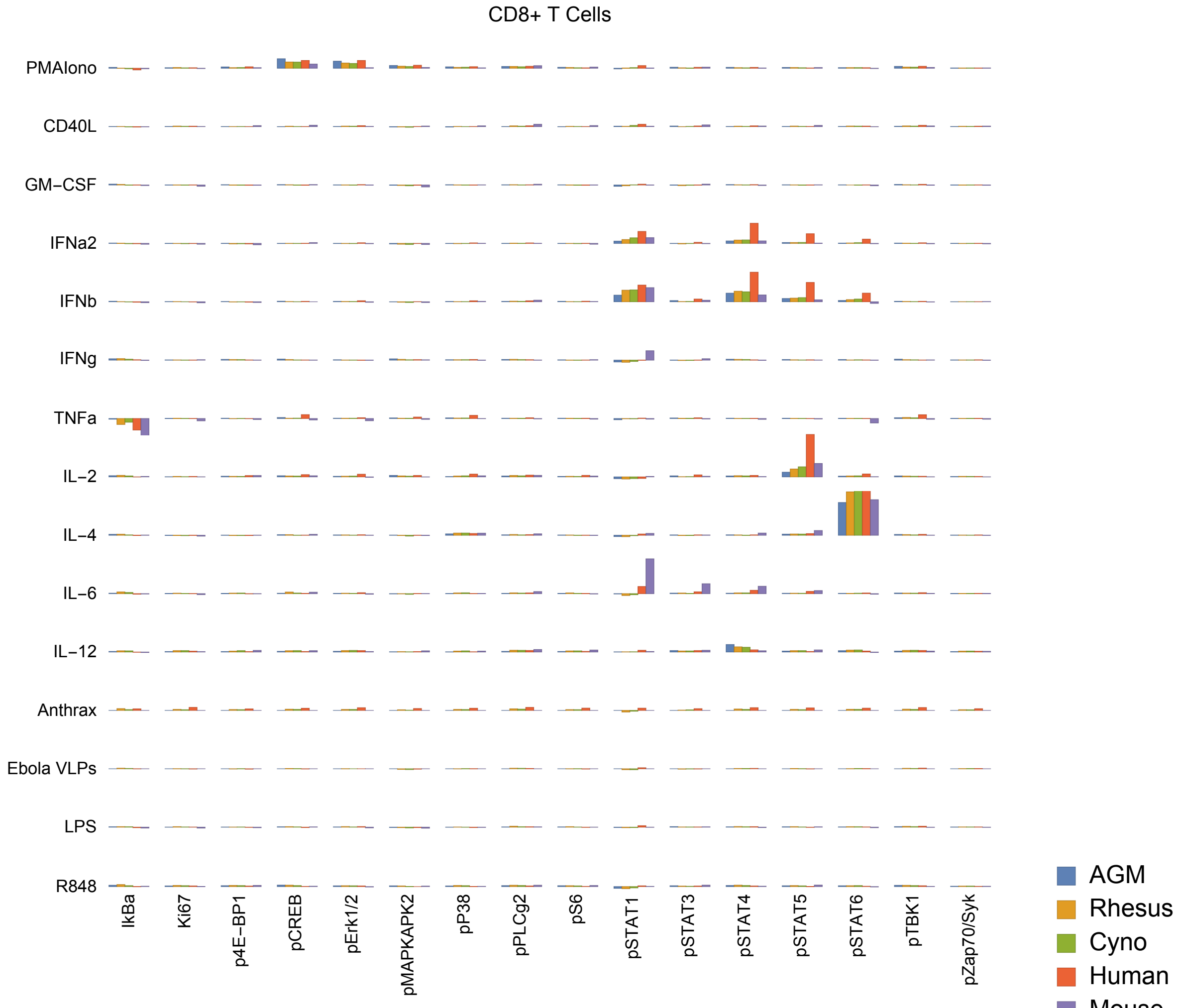
CD8+ T Cells
PMAIono
CD40L
GM−CSF
IFNa2
IFNb
IFNg
TNFa
IL−2
IL−4
IL−6
IL−12
Anthrax
Ebola VLPs
LPS
R848
IkBa
Ki67
p4E−BP1
pCREB
pErk1/2
pMAPKAPK2
pP38
pPLCg2
pS6
pSTAT1
pSTAT3
pSTAT4
pSTAT5
pSTAT6
pTBK1
pZap70/Syk
AGM
Rhesus
Cyno
Human
Mouse

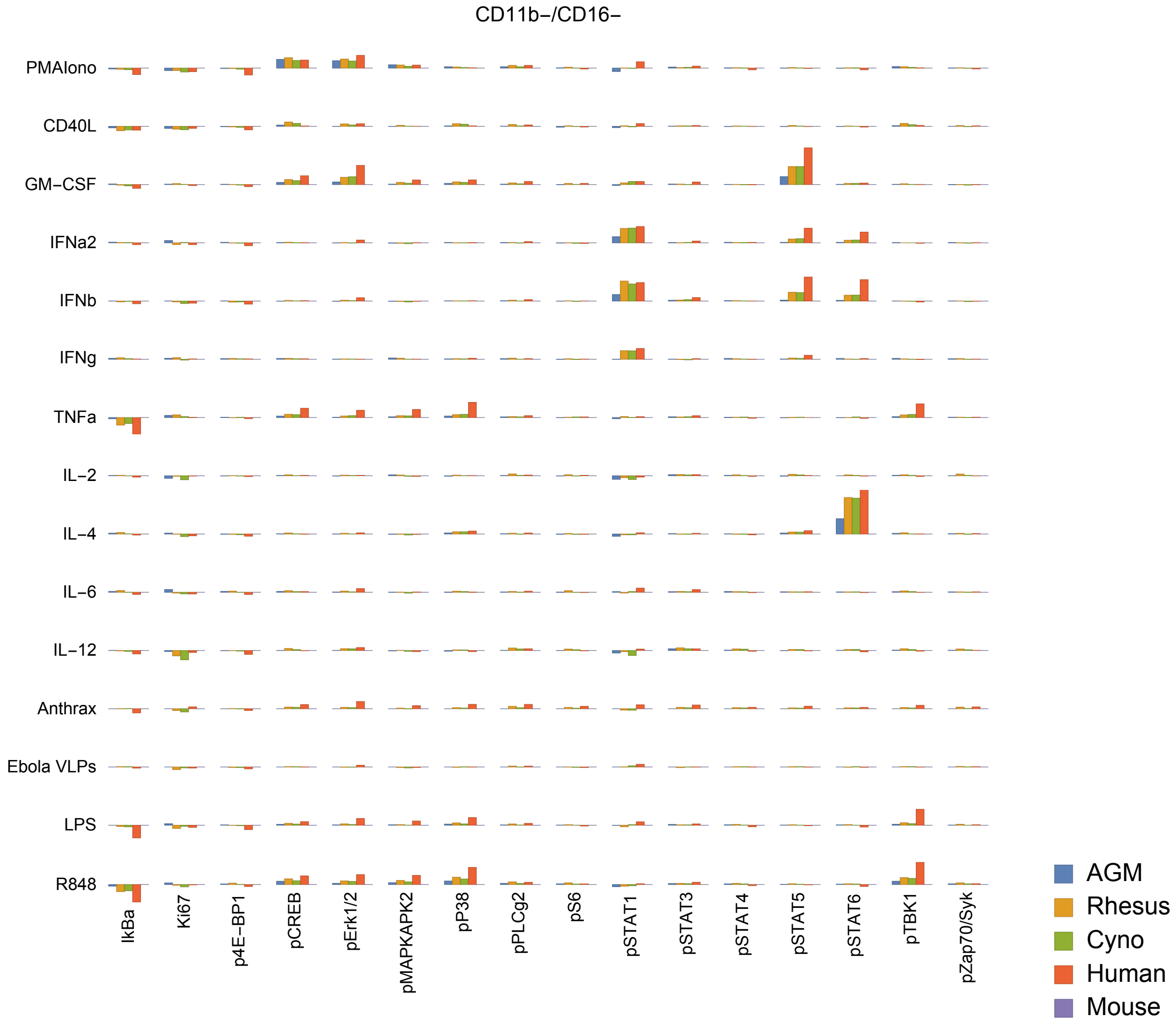
CD11b−/CD16−
PMAIono
CD40L
GM−CSF
IFNa2
IFNb
IFNg
TNFa
IL−2
IL−4
IL−6
IL−12
Anthrax
Ebola VLPs
LPS
R848
IkBa
Ki67
p4E−BP1
pCREB
pErk1/2
pMAPKAPK2
pP38
pPLCg2
pS6
pSTAT1
pSTAT3
pSTAT4
pSTAT5
pSTAT6
pTBK1
pZap70/Syk
AGM
Rhesus
Cyno
Human
Mouse

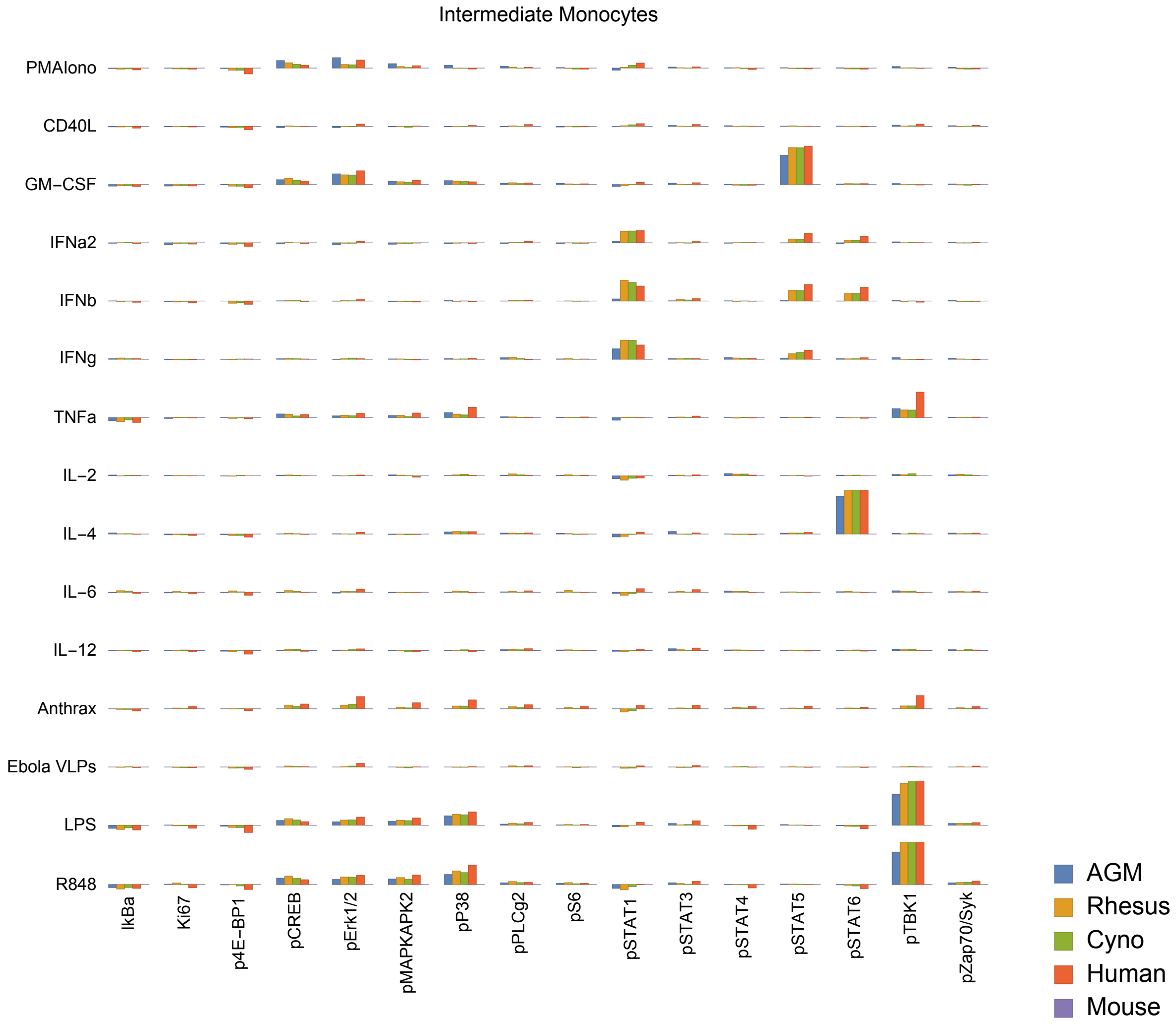
Intermediate Monocytes
PMAIono
CD40L
GM−CSF
IFNa2
IFNb
IFNg
TNFa
IL−2
IL−4
IL−6
IL−12
Anthrax
Ebola VLPs
LPS
R848
IkBa
Ki67
p4E−BP1
pCREB
pErk1/2
pMAPKAPK2
pP38
pPLCg2
pS6
pSTAT1
pSTAT3
pSTAT4
pSTAT5
pSTAT6
pTBK1
pZap70/Syk
AGM
Rhesus
Cyno
Human
Mouse

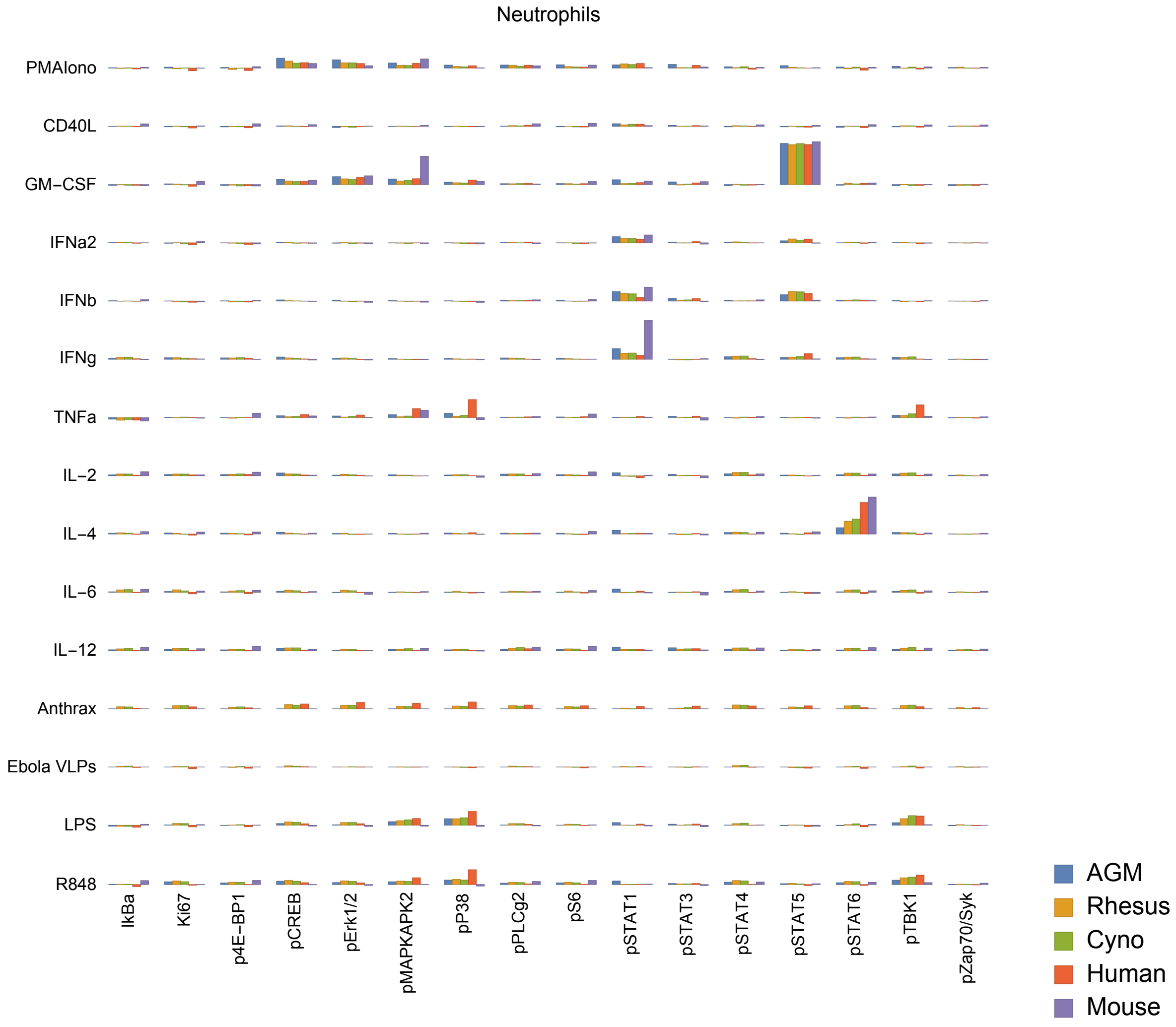

Neutrophils
PMAIono
CD40L
GM−CSF
IFNa2
IFNb
IFNg
TNFa
IL−2
IL−4
IL−6
IL−12
Anthrax
Ebola VLPs
LPS
R848
IkBa
Ki67
p4E−BP1
pCREB
pErk1/2
pMAPKAPK2
pP38
pPLCg2
pS6
pSTAT1
pSTAT3
pSTAT4
pSTAT5
pSTAT6
pTBK1
pZap70/Syk
AGM
Rhesus
Cyno
Human
Mouse

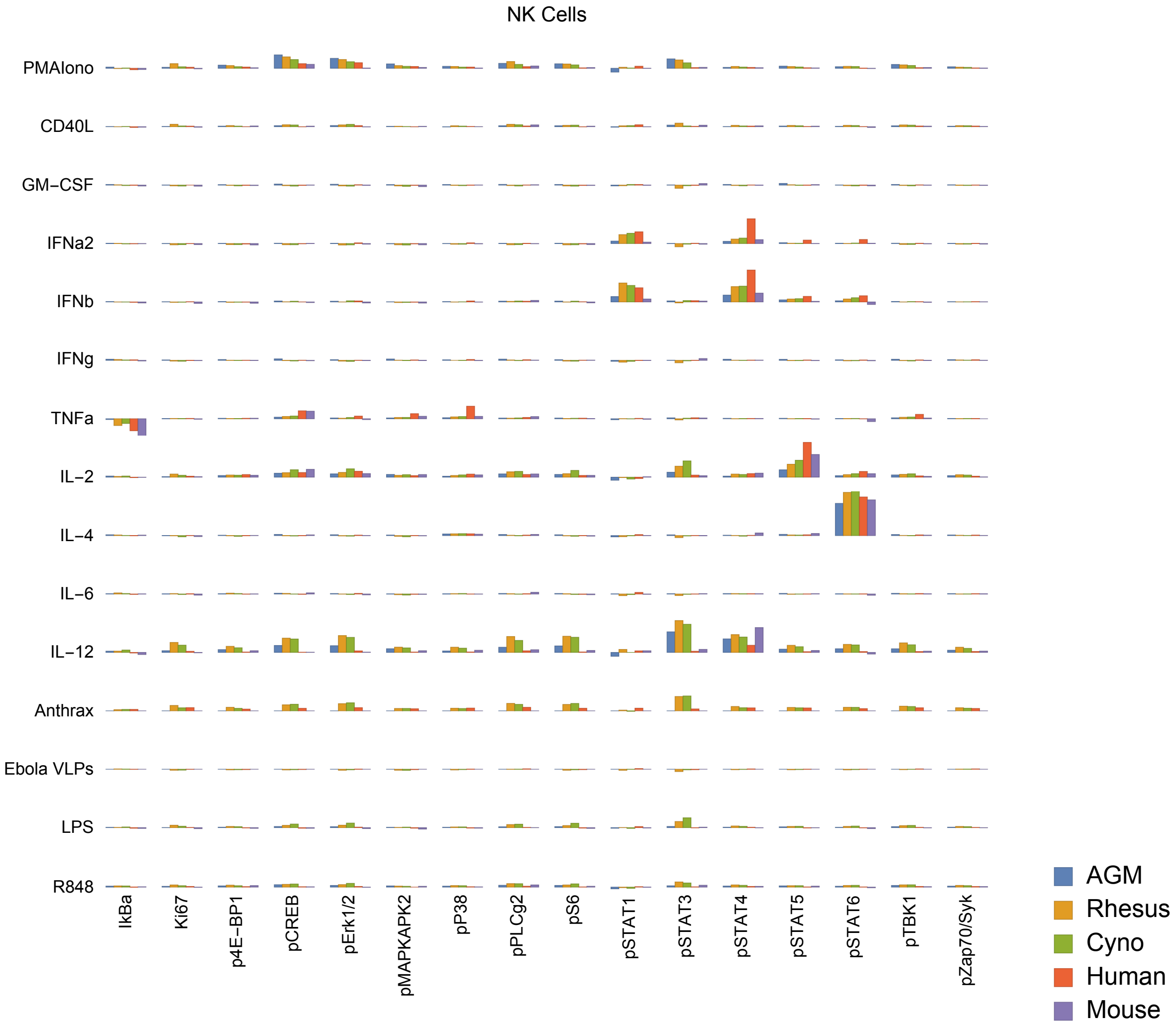
NK Cells
PMAIono
CD40L
GM−CSF
IFNa2
IFNb
IFNg
TNFa
IL−2
IL−4
IL−6
IL−12
Anthrax
Ebola VLPs
LPS
R848
IkBa
Ki67
p4E−BP1
pCREB
pErk1/2
pMAPKAPK2
pP38
pPLCg2
pS6
pSTAT1
pSTAT3
pSTAT4
pSTAT5
pSTAT6
pTBK1
pZap70/Syk
AGM
Rhesus
Cyno
Human
Mouse

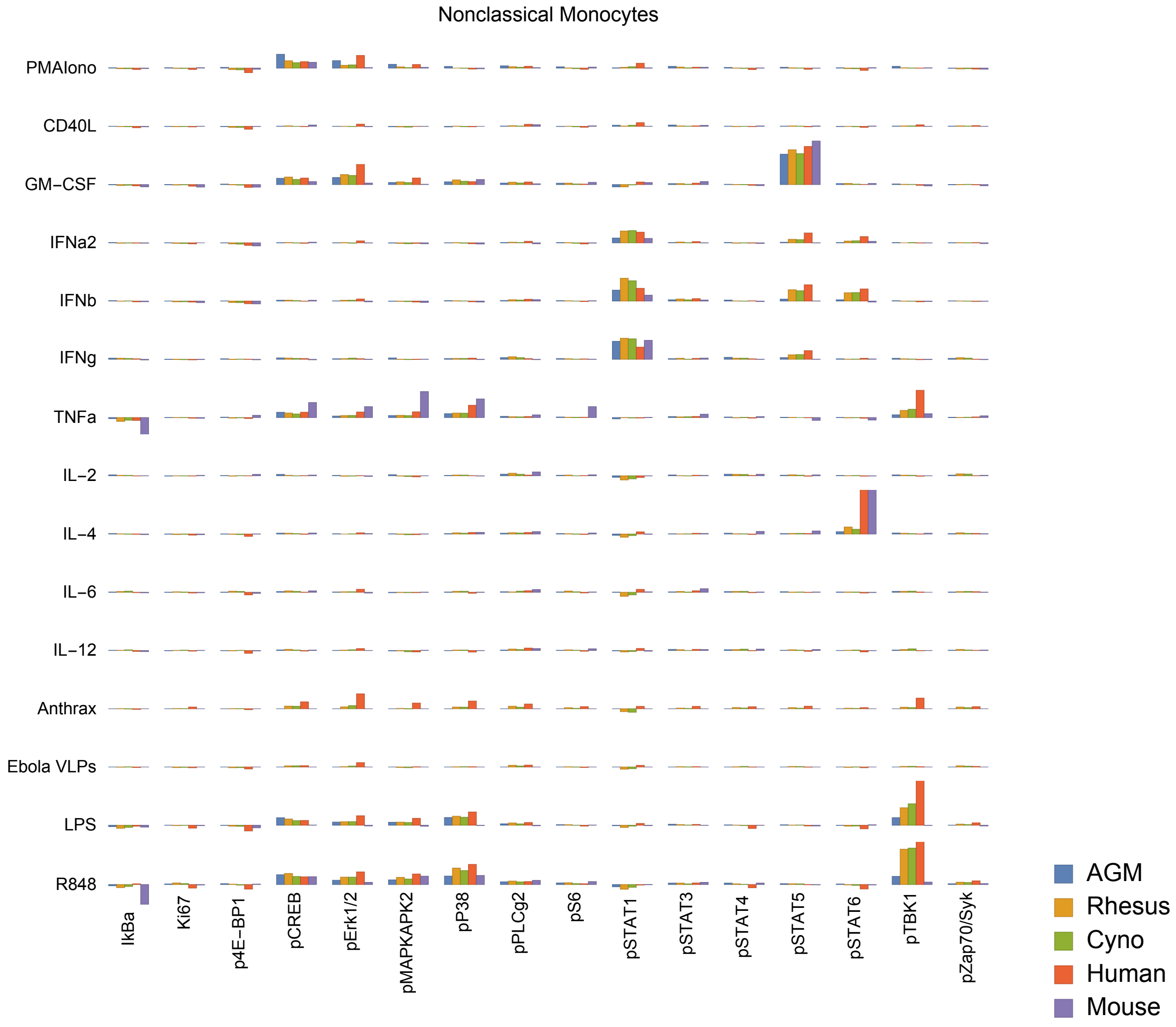

Nonclassical Monocytes
PMAIono
CD40L
GM−CSF
IFNa2
IFNb
IFNg
TNFa
IL−2
IL−4
IL−6
IL−12
Anthrax
Ebola VLPs
LPS
R848
IkBa
Ki67
p4E−BP1
pCREB
pErk1/2
pMAPKAPK2
pP38
pPLCg2
pS6
pSTAT1
pSTAT3
pSTAT4
pSTAT5
pSTAT6
pTBK1
pZap70/Syk
AGM
Rhesus
Cyno
Human
Mouse

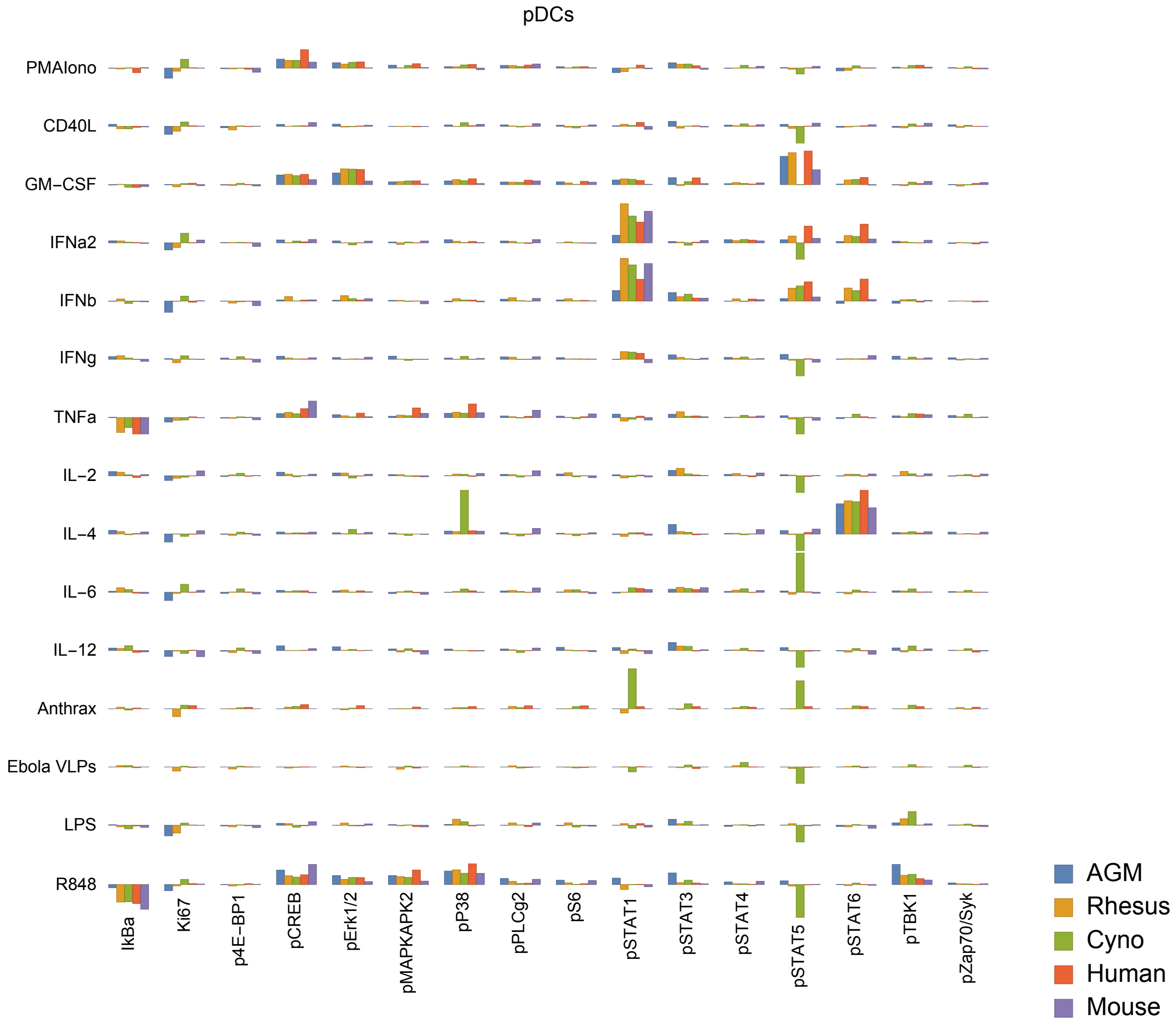
pDCs
PMAIono
CD40L
GM−CSF
IFNa2
IFNb
IFNg
TNFa
IL−2
IL−4
IL−6
IL−12
Anthrax
Ebola VLPs
LPS
R848
IkBa
Ki67
p4E−BP1
pCREB
pErk1/2
pMAPKAPK2
pP38
pPLCg2
pS6
pSTAT1
pSTAT3
pSTAT4
pSTAT5
pSTAT6
pTBK1
pZap70/Syk
AGM
Rhesus
Cyno
Human
Mouse

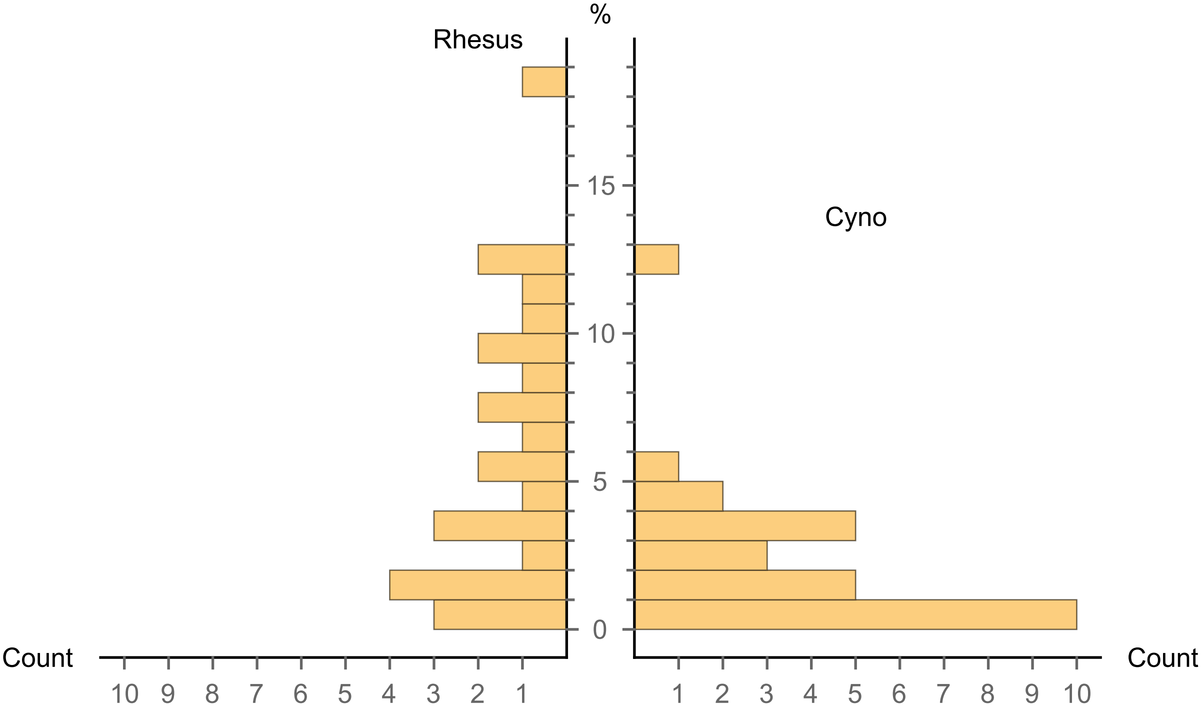


CD4+/CD8+ T cells (% of Singlet Cells)

| Species | Percent |
|---|---|
| Human | 0.110735 |
| Rhesus | 5.26584 |
| Cyno | 1.42308 |
| AGM | 0.111882 |
| Mouse | 0.183403 |

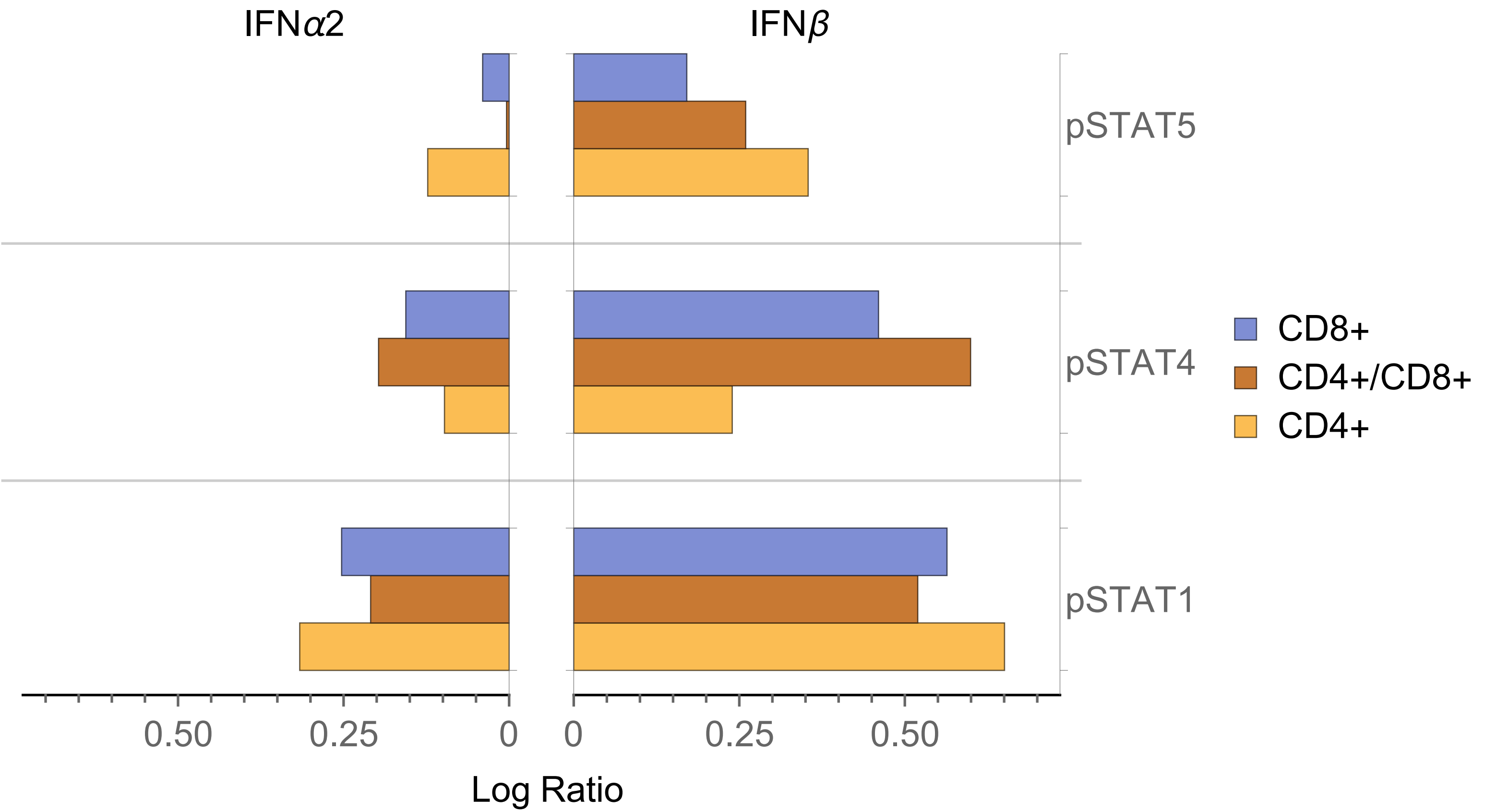
IFNα2
IFNβ
pSTAT5
pSTAT4
pSTAT1
CD8+
CD4+/CD8+
CD4+
0.50
0.25
0
0
0.25
0.50
Log Ratio

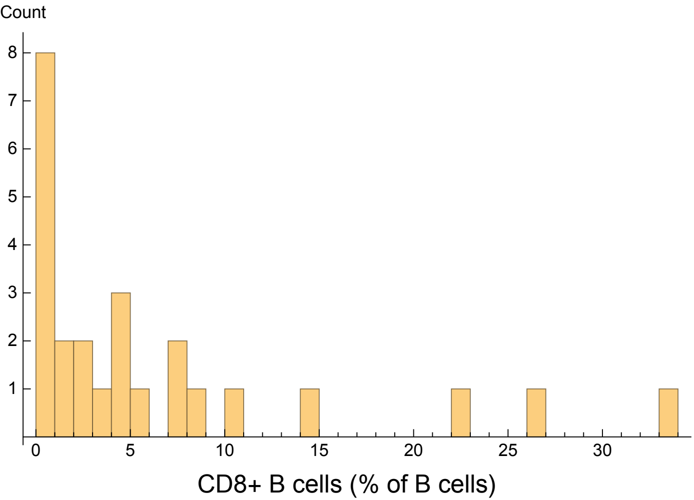
Count
8
7
6
5
4
3
2
1
0
5
10
15
20
25
30
CD8+ B cells (% of B cells)

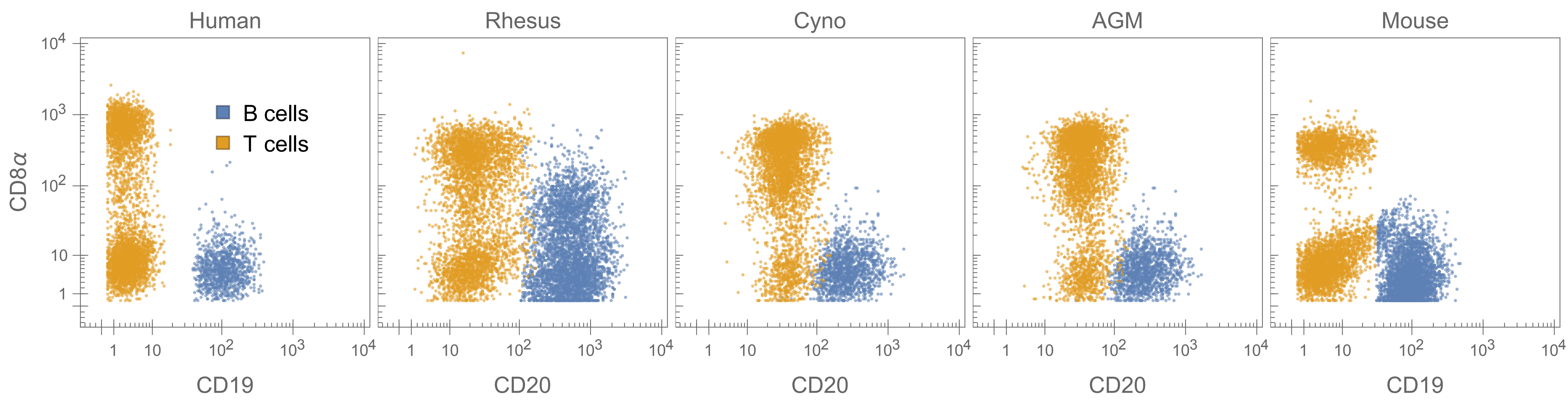
Human
Rhesus
Cyno
AGM
Mouse
CD8α
B cells
T cells
CD19
CD20
CD20
CD20
CD19

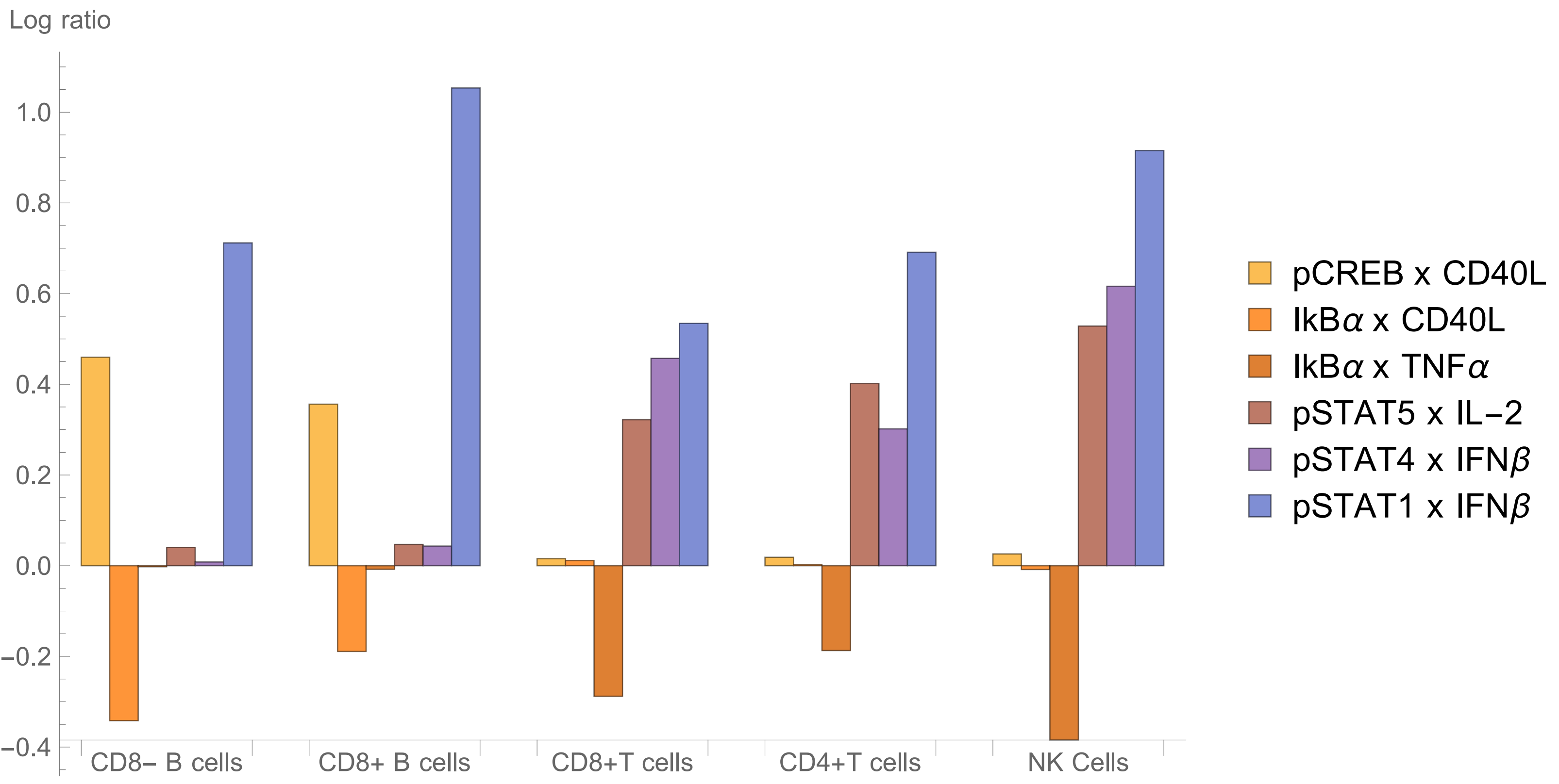

Log ratio
1.0
0.8
0.6
0.4
0.2
0.0
−0.2
−0.4
CD8− B cells
CD8+ B cells
CD8+T cells
CD4+T cells
NK Cells
pCREB x CD40L
IkBα x CD40L
IkBα x TNFα
pSTAT5 x IL−2
pSTAT4 x IFNβ
pSTAT1 x IFNβ

| Canonical Target | Human | Mouse | Macaque | African green |
|---|---|---|---|---|
| Lymph/mono | CD45 (HI30) | CD45.2 (104) and pan-CD45 (30-F11) | CD45 (DO58-1283) | CD45 (DO58-1283) |
| Erythrocytes | CD235a (HIR2) | Ter119 (TER-119) | CD233 (BRIC6) | |
| Platelets | CD61 (VI-PL2) | CD61 (2C9.G2) | CD61 (VI-PL2) | CD61 (VI-PL2) |
| Granulocytes | CD66abce (TET2) | Ly6G (1A8) | CD66abce (TET2) | CD66abce (TET2) |
| Granulocytes | | FcεRIα (MAR-1) | CD33 (AC104.3E3) | |
| Basophils | CD123 (7G3) | FcεRIα (MAR-1) | CD123 (7G3) | |
| B cells | CD19 (J3-119) | CD19 (6D5) | CD20 (2H7) | CD20 (2H7) |
| B cells | | B220 (RA3-6B2) | | |
| B cells, class-switched | IgM (G20-127) | IgM (RMM-1) | IgM (G20-127) | IgM (G20-127) |
| APCs (B, DC) | HLA-DR (Immu357) | MHC II (M5/114.15.2) | HLA-DR (Immu357) | HLA-DR (Immu357) |
| DCs, B cells | CD1c (AD5-8E7) | | CD1c (AD5-8E7) | CD1c (AD5-8E7) |
| cDCs | BDCA3 (1A4) | (CD8) | BDCA3 (1A4) | BDCA3 (1A4) |
| pDCs | CD123 (7G3) | BST-2 (120g8) | CD123 (7G3) | CD123 (7G3) |
| T subset | CCR7 (150503) | CD62L (MEL-14) | CCR7 (150503-FITC) | |
| T cells | CD3 (SP34.2) | CD3 (17A2) | CD3 (SP34.2) | CD3 (SP34.2) |
| CD4+ T cells | CD4 (OKT4) | CD4 (RM4-5) | CD4 (OKT4) | CD4 (L200) |
| CD8+ T cells | CD8 (RPA-T8) | CD8 (53-6.7) | CD8 (RPA-T8) | CD8 (RPA-T8) |
| Naïve T cells | CD45RA (HI100) | CD44 (IM7) | CD45RA (HI100) | |
| Myeloid cells | CD11b (ICRF44) | CD11b (M1/70) | CD11b (ICRF44) | CD11b (ICRF44) |
| Myeloid, T subset | CD11c (Bu15) | CD11c (HL3) | CD11c (3.9) | CD11c (3.9) |
| Monocytes | CD14 (M5E2) | Ly6C (HK1.4) | CD14 (M5E2) | CD14 (M5E2) |
| NK, mono, gran | CD16 (3G8) | CD16/32 (2.4G2) | CD16 (3G8) | CD16 (3G8) |
| Myeloid | CD33 (AC104.3E3) | CD115 (CSF-1R) | | |
| Monocytes | | | CD56 (NCAM16.2) | CD56 (NCAM16.2) |
| NK cells | CD56 (NCAM16.2) | CD49b (HMa2) | | |
| NK cells | CD7 (M-T701) | | CD7 (M-T701) | CD7 (M-T701) |
| NK cells | CD161 (HP-G310) | NK1.1/CD161c (PK126) | CD161 (HP-G310) | CD161 (HP-G310) |

| Study | Species | Origin | Age | % PBL |
|---|---|---|---|---|
| This | Rhesus | China | 5-9 years | 5.3 |
| This | Cyno | China | 5-9 years | 1.4 |
| Akari 1997[48] | Cyno | Indonesia, Philippines, Malaysia | 0-4 years<br>5-9 years<br>10-14 years<br>>15 years | 1.2<br>2.7<br>9.4<br>8.7 |
| Lee[55] | Cyno | Not specified | 6.5 years<br>7.3 years<br>7.7 years<br>8.8 years<br>9.4 years<br>10.0 years<br>11.0 years | 3.1†<br>5.1<br>4.5<br>4.7<br>7.1<br>5.0<br>6.0 |
| Reimann 1994[56] | Rhesus | Not specified | Not specified | <10 |
| Dean 1996[57] | Rhesus | Not specified | Not specified | 5.8* |
| Wang 2008[58] | Rhesus | India | 0 to 3 days<br>12 to 21 days | ~2<br>~4 |

| Stimulus | Host Species | Used In | Produced In | Lot | Working Concentration |
|---|---|---|---|---|---|
| GM-CSF | Rhesus | Rhesus Cyno | E. coli | 3600102 | 100 ng/ml |
| GM-CSF | Human | Human | E. coli | 3112711 | 100 ng/ml |
| GM-CSF | Mouse | Mouse | E. coli | 3201617 | 100 ng/ml |
| IFNα2 | Rhesus/Cyno (identical) | Rhesus Cyno | E. coli | 5967, 5616 pooled | 150 ng/ml |
| IFNα2 | Human | Human | E. coli | 5962 | 150 ng/ml |
| IFNα2 | Mouse | Mouse | E. coli | E05729-1634 | 150 ng/ml |
| LPS | N/A | Human Rhesus Cyno Mouse | E. coli O111:B4 | LEB-36-01 | 1 μg/ml |
| IL-6 | Rhesus | Rhesus Cyno | E. coli | 3600502 | 500 ng/ml |
| IL-6 | Human | Human | E. coli | 3103810 | 500 ng/ml |
| IL-6 | Mouse | Mouse | E. coli | 3201609 | 500 ng/ml |
| Resiquimod (R848) | N/A | Human Rhesus Cyno Mouse | N/A | 848-35-14 | 10 μg/ml |
| IFNγ | Rhesus | Rhesus Cyno | E. coli | ETZ0213071 | 330 ng/ml |
| IFNγ | Human | Human | E. coli | RAX1814011 | 330 ng/ml |
| IFNγ | Mouse | Mouse | E. coli | 061398 | 330 ng/ml |
| TNFα | Rhesus | Rhesus Cyno | E. coli | DCSD0113081 | 100 ng/ml |
| TNFα | Human | Human | E. coli | DDHB0113062 | 100 ng/ml |
| TNFα | Mouse | Mouse | E. coli | CS1313081 | 100 ng/ml |
| IFNβ* | Human | Human Rhesus Cyno | CHO | 5886 | 5 ng/ml |
| IFNβ | Mouse | Mouse | Human cell line | Missing | |
| CD40L soluble dimer (“MegaCD40L”) | Human | Human Rhesus Cyno | CHO | 05281412, 03041401 pooled | 125 ng/ml |
| CD40L soluble dimer (“MegaCD40L”) | Mouse | Mouse | CHO | 01151321 | 125 ng/ml |
| PMA and ionomycin* | N/A | Human Rhesus Cyno Mouse | N/A | E13495-116 | 0.081 μM PMA and 1.34 μM iono |
| IL-12 | Rhesus | Rhesus Cyno | CHO | OQM0210071 | 400 ng/ml |
| IL-12 | Human | Human | CHO | 0210596, 0707596-2 pooled | 400 ng/ml |
| IL-12 | Mouse | Mouse | CHO | 0407S97 | 400 ng/ml |
| IL-4 | Rhesus | Rhesus Cyno | E. coli | IXV0113111 | 125 ng/ml |
| IL-4 | Human | Human | E. coli | AG1314021 | 125 ng/ml |
| IL-4 | Mouse | Mouse | E. coli | BC1613061 | 125 ng/ml |

Commented [b1]:

| Stimulus | Host Species | Used In | Produced In | Lot | Working Concentration |
|---|---|---|---|---|---|
| IL-2 | Rhesus/Cyno (identical) | Rhesus Cyno | E. coli | 04/12/2008 | 2 µg/ml |
| IL-2 | Human | Human | E. coli | 101312, 041412 pooled | 2 µg/ml |
| IL-2 | Mouse | Mouse | E. coli | 0608108 | 2 µg/ml |
| *Gamma-inactivated vegetative *Bacillus anthracis* Ames | N/A | Human Rhesus Cyno | | AGD0001331 | 400,000 CFU/ml |
| *Zaïre Ebolavirus-like particles | N/A | Human Rhesus Cyno | 293T cells | | Varied |

| Antigen | Clone | Label | Vendor and Catalog | Staining Concentration (µg/ml) |
|---|---|---|---|---|
| CD45 | HI30 | 115 | Biolegend 304002 | 3.25 |
| CD235a | HIR2 | 113 | Biolegend 306602 | 0.67 |
| CD61 | VI-PL2 | 140 | Biolegend 336402 | 5.37 |
| CD66 | YTH71.3 | 158 | Pierce MA1-36189 | 3.41 |
| CD19 | J3-119 | 173 | Beckman Coulter IM1313 | 1.79 |
| IgM | G20-127 | 174 | BD 555780 | 2.68 |
| HLA-DR | Immu357 | 176 | Beckman Coulter Immu357 | 0.34 |
| CD1c | AD5-8E7 | 162 | Miltenyi (Custom) | 1.34 |
| BDCA3 | 1A4 | 164 | BD 559780 | 4.31 |
| CD123 | 7G3 | 148 | BD 554527 | 1.34 |
| CCR7 | 150503 | 171 | R&D MAB197-100 | 7.01 |
| CD3 | SP34.2 | 157 | BD 551916 | 2.68 |
| CD4* | OKT4 | 156 | Biolegend 317402 | 1.25 |
| CD8 | RPA-T8 | 155 | Biolegend 301002 | 1.41 |
| CD45RA | HI100 | 166 | Biolegend 304102 | 2.68 |
| CD11b | ICRF44 | 153 | Biolegend 301302 | 7.16 |
| CD11c | Bu15 | 143 | Biolegend 337202 | 1.34 |
| CD14 | M5E2 | 151 | Biolegend 301802 | 10.73 |
| CD16 | 3G8 | 159 | Biolegend 302033 | 2.68 |
| CD33 | AC104.3E3 | 142 | Miltenyi (Custom) | 3.58 |
| CD56 | NCAM16.2 | 175 | BD 559043 | 1.79 |
| CD7 | M-T701 | 141 | BD 555359 | 5.37 |
| CD161 | HP-G310 | 168 | Biolegend 339902 | 5.37 |

| Antigen | Clone | Label | Vendor and Catalog | Straining Concentration (µg/ml) |
|---|---|---|---|---|
| CD45 | DO58-1283 | 115 | BD 552566 | 5.37 |
| CD233 | BRIC 6 | 113 | IBGRL 9439 | 4.11 |
| CD61 | VI-PL2 | 140 | Biolegend 336402 | 5.37 |
| CD66 | YTH71.3 | 158 | Thermo MA5-17003 | 2.25 |
| CD20 | 2H7 | 173 | Biolegend 302302 | 5.37 |
| IgM | G20-127 | 174 | BD 555780 | 2.68 |
| HLA-DR | Immu357 | 176 | Beckman Coulter | 0.34 |
| CD1c | AD5-8E7 | 162 | Miltenyi | 1.34 |
| BDCA3 | 1A4 | 164 | BD 559780 | 3.58 |
| CD123 | 7G3 | 148 | BD 554527 | 1.34 |
| CCR7 | 150503 | FITC | BD 561271 | 8.94 |
| FITC | FIT-22 | 171 | Biolegend 408302 | 5.39 |
| CD3 | SP34.2 | 157 | BD 551916 | 2.68 |
| CD4* | OKT4 | 156 | Biolegend 317404 | 1.25 |
| CD8 | RPA-T8 | 155 | Biolegend 301002 | 1.41 |
| CD45RA | HI100 | 166 | Biolegend 304102 | 2.68 |
| CD11b | ICRF44 | 153 | Biolegend 301312 | 6.71 |
| CD11c | 3.9 | 143 | Biolegend | 1.34 |
| CD14 | M5E2 | 151 | Biolegend 301810 | 10.73 |
| CD16 | 3G8 | 159 | Biolegend 302033 | 2.68 |
| CD33 | AC104.3E3 | 142 | Miltenyi | 4.29 |
| CD56 | NCAM16.2 | 175 | BD 559403 | 1.79 |
| CD7 | M-T701 | 141 | BD 555359 | 5.37 |
| CD161 | HP-G310 | 168 | Biolegend 339902 | 3.08 |

| Antigen | Clone | Label | Vendor and Catalog | Straining Concentration (μg/ml) |
|---|---|---|---|---|
| Ter119 | TER-119 | 113 | Biolegend 116202 | 5.37 |
| CD45.2 | 104 | 115 | Biolegend 109801 | 5.37 |
| pan-CD45 | 30-F11 | 115 | Biolegend 103102 | 5.37 |
| CD61 | 2C9.G2 | 140 | Biolegend 104302 | 1.34 |
| Ly6G | 1A8 | 158 | Biolegend 127602 | 0.34 |
| CD19 | 6D5 | 173 | Biolegend 115502 | 1.34 |
| B220 | RA3-6B2 | 141 | Biolegend 103202 | 22.68 |
| IgM | RMM-1 | 174 | Biolegend 406501 | 2.68 |
| MHC II | M5/114.15.2 | 176 | Biolegend 107602 | 0.34 |
| BST-2 | 120g8 | 148 | Dendritics DDX0390P-100 | 1.34 |
| CD115 | CSF-1R | 164 | Biolegend 135502 | 1.34 |
| FcεRIa | MAR-1 | 166 | Bioelgend 134302 | 0.34 |
| CD62L | MEL-14 | FITC | Biolegend 104406 | 3.22 |
| CD3 | 17A2 | 157 | Biolegend 100202 | 0.34 |
| CD4 | RM4-5 | 156 | Biolegend 100506 | 0.34 |
| CD8 | 53-6.7 | 155 | Biolegend 100702 | 0.34 |
| CD44 | IM7 | 175 | Biolegend 103002 | 0.34 |
| CD11b | M1/70 | 153 | Biolegend 101202 | 1.34 |
| CD11c | HL3 | 143 | BD 553799 | 0.34 |
| Ly6C | HK1.4 | 151 | Biolegend 128002 | 0.34 |
| CD16/32 | 2.4G2 | 162 | BD 553142 | 1.34 |
| NK1.1 | PK136 | 168 | Biolegend 108702 | 1.34 |
| CD49b | HMa2 | 142 | Biolegend 103501 | 0.34 |

| Antigen | Label | Species | Clone | Straining Concentration (μg/ml) |
|---|---|---|---|---|
| STAT1 pY701 | 147 | primates<br>mice | 4a | 3.83<br>6.97 |
| STAT3 pY705 | 139 | all | 4 | 5.82 |
| STAT4 pY693 | 170 | all | 38 | 5.66 |
| STAT5 pY694 | 149 | all | 46 | 5.37 |
| STAT6 pY691 | 165 | primates<br>mice | 18<br>J71-773.58.11 | 5.38<br>8.33 |
| Ki67 | 169 | all | SolA15 | 3.36 |
| Erk1/2 pT202/Y204 | 152 | all | D13.14.4E | 16.44 |
| MAPKAPK2 pT334 | 144 | all | 27B7 | 2.29 |
| CREB pS133 | 145 | all | 87G3 | 7.24 |
| IkBa amino-terminal | 163 | all | L35A5 | 3.79 |
| TBK1/NAK pS172 | 161 | all | D52C2 | 10.78 |
| S6 pS235/236 | 150 | all | 2F9 | 13.15 |
| Zap70/Syk pY319/Y352 | 160 | all | 17a | 2.14 |
| 4E-BP1 pT37/46 | 172 | all | 236B4 | 3.13 |
| PLCγ2 pY759 | 146 | all | K86-689.37 | 2.18 |
| P38 pT180/Y182 | 154 | all | 36/p38 | 3.56 |
| FITC (for CCR7) | 171 | NHPs | FIT-22 | 5.33 |
| FITC (for CD62L) | 171 | mice | FIT-22 | 8.44 |
| FoxP3 | 167 | primates<br>mice | PCH101<br>NRRF-30 | 10.55<br>7.16 |
